# Charting vulnerability of brainstem functional pathways in aging and prodromal synucleinopathy

**DOI:** 10.64898/2026.10.27.760186

**Authors:** Lin Hua, Kavita Singh, María G. García-Gomar, F. Fabrice Hannanu, Simone Cauzzo, Subhranil Koley, Ambra Stefani, Stephan Grimaldi, Aleksandar Videnovic, Marta Bianciardi

**Affiliations:** Brainstem Imaging Laboratory, Department of Radiology, Athinoula A. Martinos Center for Biomedical Imaging, Massachusetts General Hospital and Harvard Medical School, Boston, MA, USA; Department of Radiology and Bioengineering, School of Medicine, Uniformed Services University of the Health Sciences, Bethesda, MD, USA; Henry M. Jackson Foundation for the Advancement of Military Medicine, Inc., Bethesda, MD, USA; Escuela Nacional de Estudios Superiores Unidad Juriquilla, Universidad Nacional Autónoma de México, Querétaro, Mexico; National Institute on Alcoholism and Alcohol Abuse, National Institutes of Health, Bethesda, MD, USA; Department of Neuroscience, University of Padova, Padova, Italy; Department of Neurology, Massachusetts General Hospital and Harvard Medical School, Boston, MA, USA; Department of Neurology, Medical University of Innsbruck, Innsbruck, Austria; Department of Neurology and Movement Disorders, APHM, Timone University Hospital, Marseille, France; CRMBM-CEMEREM, School of Medicine, Aix Marseille University, Marseille, France; Division of Sleep Medicine, Harvard University, Boston, MA, USA

**Keywords:** 7-Tesla fMRI, Human brainstem-cortical functional connectome, Brainstem, Aging, Isolated REM sleep behavior disorder (iRBD)

## Abstract

The brainstem is vulnerable to aging and prodromal synucleinopathy, yet whether these processes disrupt shared or distinct functional pathways linking the brainstem to the cortex remains unknown. Using high-resolution 7-Tesla fMRI, we mapped connectivity between 58 brainstem nuclei and cortical regions in young adults, older adults and individuals with isolated REM sleep behavior disorder (iRBD), a prodromal synucleinopathy. Long-range brainstem-cortical connectivity weakened with aging and further in iRBD. However, the two conditions affected distinct systems: aging involved arousal- and cognition-related nuclei and association cortex, whereas iRBD involved sleep- and autonomic-related nuclei and extended to sensory-motor cortex. These alterations were associated with distinct neurophysiological, metabolic and neurotransmitter architectures and spatial transcriptomic signatures. Brainstem-cortical connectivity also followed the cortical unimodal-transmodal hierarchy, with reduced differentiation in aging. Independent 3-Tesla data reproduced the broader connectivity patterns. Together, our findings reveal shared and distinct multiscale patterns of brainstem pathway vulnerability in aging and prodromal synucleinopathy.

## Introduction

The brainstem is an evolutionarily conserved structure that regulates functions essential for survival and consciousness through extensive ascending and descending pathways and widespread neuromodulatory projections connecting the spinal cord with subcortical nuclei and cortical regions, thereby shaping activity across the nervous system^<u>1–3</u>^. This central anatomical and functional position places the brainstem at a critical intersection in multiple brain diseases, including disorders of consciousness, sleep disorders, dysautonomia, mental illness, and neurodegenerative diseases, and makes its connectivity a potentially important substrate of disease-related functional alterations^<u>4–10</u>^.

Accumulating evidence from animal models and brainstem lesion studies indicates that brainstem nuclei undergo selective, structural and functionally meaningful alterations during aging, some of which may overlap with pathways vulnerable in neurodegenerative diseases such as synucleinopathy. For instance, noradrenergic neurons and serotonergic projections from the locus coeruleus and raphe nuclei show structural and functional abnormalities in pathways involving the prefrontal cortex and hippocampus in aged rodents, accompanied by impairments in cognitive and motor function^<u>11–14</u>^. In the context of synucleinopathy, isolated rapid eye movement sleep (REM) behavior disorder (iRBD), characterized by the loss of muscle atonia during REM sleep and the emergence of dream enactment behaviors^<u>15</u>^, is closely linked to dysfunction of brainstem circuits controlling REM sleep atonia^<u>5</u>^ and is recognized as one of the earliest clinical manifestations of synucleinopathy^<u>16,17</u>^. Classical lesion studies implicate the medullary magnocellular reticular formation, locus coeruleus/subcoeruleus complex, pedunculopontine nucleus, laterodorsal tegmental nucleus, and possibly substantia nigra as structures relevant to iRBD pathophysiology^<u>5</u>^. Subsequent rodent studies identify distinct REM-off regions in the ventrolateral periaqueductal gray and lateral pontine tegmentum and REM-on regions including the sublaterodorsal nucleus and precoeruleus^<u>18</u>^. Lesions of the sublaterodorsal nucleus disrupt glutamatergic projections to inhibitory spinal interneurons, resulting in disinhibition of motor neurons and loss of REM sleep atonia^<u>18–20</u>^. Together, these findings provide consistent evidence implicating discrete brainstem nuclei in both age-related functional decline and prodromal synucleinopathy pathophysiology, while raising the question of whether aging and prodromal synucleinopathy affect shared or distinct brainstem pathways at the whole-brain level.

Despite the critical involvement of the brainstem in aging and prodromal synucleinopathy, *in-vivo* human evidence for brainstem functional alterations has largely been derived from studies focused on individual brainstem nuclei or restricted brainstem-cortical circuits. Recent resting-state fMRI evidence demonstrates age-related reductions in functional connectivity (FC) involving nuclei of the ascending arousal system and cortical association networks, particularly the default mode and salience networks^<u>21</u>^. Seed-based FC studies further reveal altered connectivity of the locus coeruleus and ventral tegmental area/substantia nigra pars compacta with cerebellar, frontal, visual and somatosensory cortical regions, suggesting age-related alterations in catecholaminergic circuits supporting cognitive and motor control^<u>22</u>^. Complementary structural connectivity findings indicate age-related degeneration of the mesolimbic tract connecting the ventral tegmental area to the nucleus accumbens, potentially contributing to decline in memory and social interaction with aging^<u>23</u>^. Evidence of brainstem FC alterations has also emerged in prodromal synucleinopathy. Reduced FC in iRBD has been reported between the substantia nigra and the ipsilateral putamen and superior occipital gyrus^<u>24</u>^, as well as between the midbrain and pallidum within the basal ganglia network^<u>25</u>^. Altered locus coeruleus FC has also been found across the default mode, executive control, salience, sensorimotor and visual networks^<u>26</u>^. Furthermore, structural connectivity studies in iRBD reveal impaired connectivity across 14 brainstem nuclei, including connections between REM-on (e.g., subcoeruleus) and REM sleep muscle atonia (e.g., medullary reticular formation) areas^<u>27</u>^. However, these studies provide a fragmented view of brainstem involvement, centered on selected nuclei and pathways. Systematic characterization has been hindered by the small size of brainstem nuclei, limited imaging resolution and sensitivity, and the historical lack of comprehensive *in-vivo* probabilistic atlases of human brainstem nuclei. Consequently, brainstem alterations have largely been examined nucleus-by-nucleus, leaving unresolved how aging and prodromal synucleinopathy reshape brainstem functional architecture as a whole, whether specific short-range cortico-cortical or long-range brainstem-cortical pathways are preferentially affected, and how these alterations relate to the brain’s broader functional hierarchy and underlying neurobiology.

In the present study, we addressed these limitations by leveraging high-sensitivity and high-spatial-resolution 7-Tesla resting-state fMRI together with a recently developed *in-vivo* probabilistic atlas encompassing 58 human brainstem nuclei to systematically investigate alterations in brainstem functional architecture associated with aging and iRBD. We asked whether aging and prodromal synucleinopathy exhibit shared or distinct patterns of brainstem pathway vulnerability, and whether these patterns can be situated within the cortical functional hierarchy and linked to underlying neurobiological and molecular architectures. First, we mapped the spatial organization of FC within and between the brainstem and cortex, testing whether vulnerability varies with connection distance across aging and iRBD. Next, we identified hubs of altered brainstem-cortical connectivity and characterized the relationship between altered connectivity and neurophysiological, structural, metabolic and chemoarchitectural features in aging and iRBD. We then examined whether the cortical functional hierarchy delineating unimodal (lower-order) and transmodal (higher-order) brain regions is reflected in brainstem connectivity patterns and whether its organization (e.g., network segregation) is altered in aging and iRBD. Finally, we integrated the spatial transcriptomic data to identify molecular signatures associated with brainstem alterations in aging and iRBD. Collectively, this multiscale framework enabled us to chart brainstem functional pathway vulnerability across aging and prodromal synucleinopathy, linking macroscale functional reorganization to cortical organization, as well as neurobiological and molecular architectures.

## Results

Resting-state fMRI time series in the brainstem and cortex were acquired on a 7-Tesla scanner from 20 younger adults (YA, age: 29.35 ± 5.4 years; 10 females), 17 older adults (OA, age: 64.41 ± 4.15 years; 3 females), and 16 individuals with iRBD (age: 67.19 ± 5.96 years; 3 females), and 3-Tesla replication data were obtained from 30 OA (age: 67.14 ± 11.83 years; 11 females) and 30 individuals with iRBD (age: 69.06 ± 5.13 years; 6 females) from the Parkinson’s Progression Markers Initiative (PPMI) dataset. Functional images were processed following established brainstem-specific protocols (Figure S1; see Method for details), and all functional connections were defined based on specified cortical and brainstem regions. Cortical regions were defined according to the 400 regions in the Schaefer parcellation^<u>28</u>^, and brainstem nuclei were defined according to the 58 nuclei in the Brainstem Navigator atlas (8 midline and 25 bilateral nuclei; Figure 1A and Figure S2).

**Figure 1.**
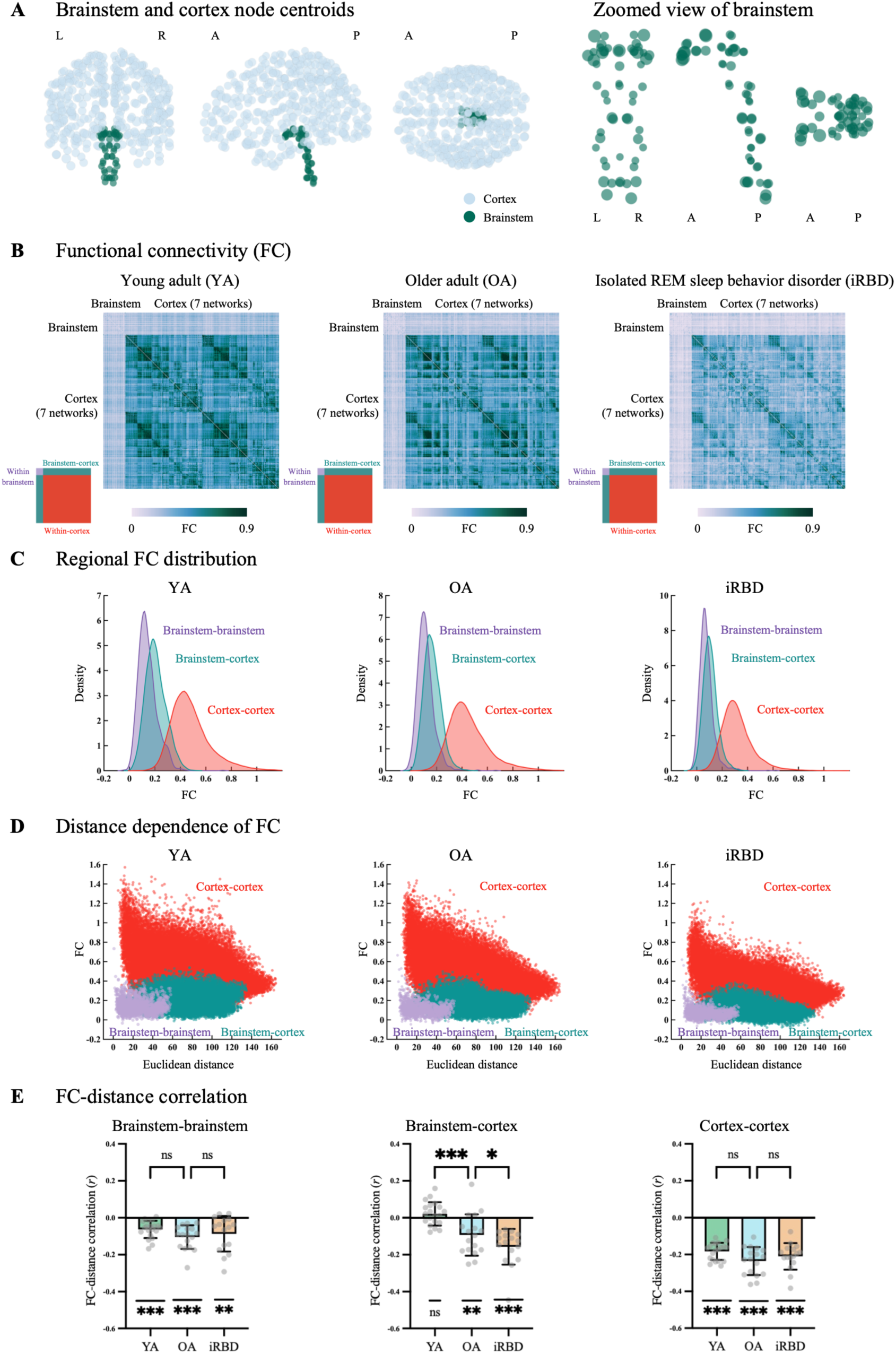
Altered distance-dependent organization of brainstem networks in aging and iRBD. (A) Parcel coordinate centroids for 58 brainstem nuclei (green) and 400 cortical regions (grey) displayed in coronal, sagittal and horizontal views. (B) Group-average whole-brain FC matrices for YA, OA and iRBD. Values represent Fisher z-transformed correlation coefficients, with the color bar denoting FC strength. Insets indicate the FC matrix regions corresponding to within-brainstem, brainstem-cortex and within-cortex connectivity. (C) Density distributions of regional FC within the brainstem (YA: 0.14 ± 0.04; OA: 0.11 ± 0.04; iRBD: 0.07 ± 0.04; purple), between brainstem and cortex (YA: 0.21 ± 0.06; OA: 0.16 ± 0.06; iRBD: 0.11 ± 0.06; green), and within the cortex (YA: 0.48 ± 0.13; OA: 0.45 ± 0.12; iRBD: 0.32 ± 0.14; red). (D) Scatter plots of the relationship between regional FC strength and Euclidean distance across groups. (E) Regional FC-distance correlation coefficients across groups: within the brainstem (one-sample *t*-tests against zero; YA: *t*_(19)_ = 5.99, *p* < 0.001; OA: *t*_(16)_ = 6.73, *p* < 0.001; iRBD: *t*_(15)_ = 3.6, *p* = 0.003), between brainstem and cortex (YA: *t*_(19)_ = 1.5, *p* = 0.15; OA: *t*_(16)_ = 3.42, *p* = 0.003; iRBD: *t*_(15)_ = 6.5, *p* < 0.001), and within the cortex: (YA: *t*_(19)_ = 17.5, *p* < 0.001; OA: *t*_(16)_ = 12.8, *p* < 0.001; iRBD: *t*_(15)_ = 11.7, *p* < 0.001). Error bars represent the standard deviation (SD). * *p* < 0.05, ** *p* < 0.01, *** *p* < 0.001, *ns* not significant.

### Long-range brainstem-cortical connectivity alterations

Regionally averaged FC strength progressively declined from YA to OA, and further to iRBD (one-way ANOVA; within-brainstem: *F*_(2,50)_ = 14.26, *p* < 0.001; brainstem-cortex: *F*_(2,50)_ = 12.44, *p* < 0.001; within-cortex: *F*_(2,50)_ = 7.39, *p* = 0.002; Figure 1B). Notably, the brainstem was more functionally connected with the cortex than with itself across all groups (paired-sample *t*-tests; YA: *t*_(19)_ = 6.07, *p* < 0.001; OA: *t*_(16)_ = 5.71, *p* < 0.001; iRBD: *t*_(15)_ = 2.15, *p* = 0.048). Consistently, the distributions of regional FC shifted toward lower values from YA to OA and iRBD for within-brainstem, brainstem-cortical and within-cortex FC (Figure 1C). Indeed, whereas FC significantly decreased with increasing Euclidean distance both within the brainstem and within the cortex, with comparable correlations across groups (one-way ANOVA; within-brainstem: *F*_(2,50)_ = 1.68, *p* = 0.197; within-cortex: *F*_(2,50)_ = 3.01, *p* = 0.058), brainstem-cortical FC showed a distinct distance-dependent pattern (one-way ANOVA; *F*_(2,50)_ = 17.8, *p* < 0.001; Figure 1D and 1E). Long-range brainstem-cortical FC remained relatively insensitive to increasing distance in YA, but decreased with increasing distance in OA and iRBD. The progressively stronger distance dependence suggests disrupted long-range information transfer from the brainstem to the cortex in aging (FDR-corrected *p* = 0.001) and iRBD (FDR-corrected *p* = 0.048).

### Brainstem-cortical alterations reflect distinct neurobiological profiles

The overall patterns of brainstem-cortical connectivity strength were further quantified using FC weighted degree to capture the global integrative capacity of individual regions. Brainstem-to-cortical weighted degree was defined as the sum of each brainstem nucleus’s FC and all cortical regions (Figure 2A). Group comparisons revealed significant age-related reductions in brainstem-to-cortical weighted degree, affecting midbrain nuclei (e.g., left cuneiform nucleus, left isthmic reticular formation, dorsal raphe, and periaqueductal gray), pontine nuclei (e.g., left lateral and bilateral medial parabrachial nuclei, bilateral pontine reticular nucleus, oral and caudal parts, left vestibular nuclei complex, right subcoeruleus, and right laterodorsal tegmental nucleus-central gray of the rhomboencephalon), and medullary nuclei (e.g., bilateral viscero-sensory-motor nuclei complex, left parvicellular reticular nucleus, alpha part, left superior and left inferior medullary reticular formations, right inferior olivary nucleus, and raphe pallidus; FDR-corrected *ps* < 0.05; Figure 2B). Functional profiling indicated predominant contributions from arousal (39.13%, *p_perm_* = 0.0147) and cognitive functions (20.14%, *p_perm_* = 0.043; Figure 3A). iRBD-related reductions exhibited distinct profiles, involving midbrain nuclei (e.g., bilateral inferior colliculus, left substantia nigra, left ventral tegmental area-parabrachial pigmented nucleus complex, and right mesencephalic reticular formation), pontine nuclei (e.g., bilateral laterodorsal tegmental nucleus-central gray of the rhomboencephalon, bilateral pontine reticular nucleus, oral and caudal parts, bilateral vestibular nuclei complex, left subcoeruleus, left medial and right lateral parabrachial nuclei, right superior olivary complex, and right locus coeruleus), and medullary nuclei (e.g., bilateral superior medullary reticular formation, left inferior olivary nucleus, and right inferior medullary reticular formation; FDR-corrected *ps* < 0.05; Figure 2B). In contrast to aging, iRBD-related alterations were predominantly associated with sleep (49.6%, *p_perm_* = 0.0007) and autonomic functions (24.3%, *p_perm_* = 0.0348; Figure 3A). Extending these findings beyond the cortex revealed progressive reductions in brainstem connectivity with specific subcortical regions in aging and iRBD. (see Supplementary Material for brainstem-subcortical connectivity; Figure S4).

**Figure 2.**
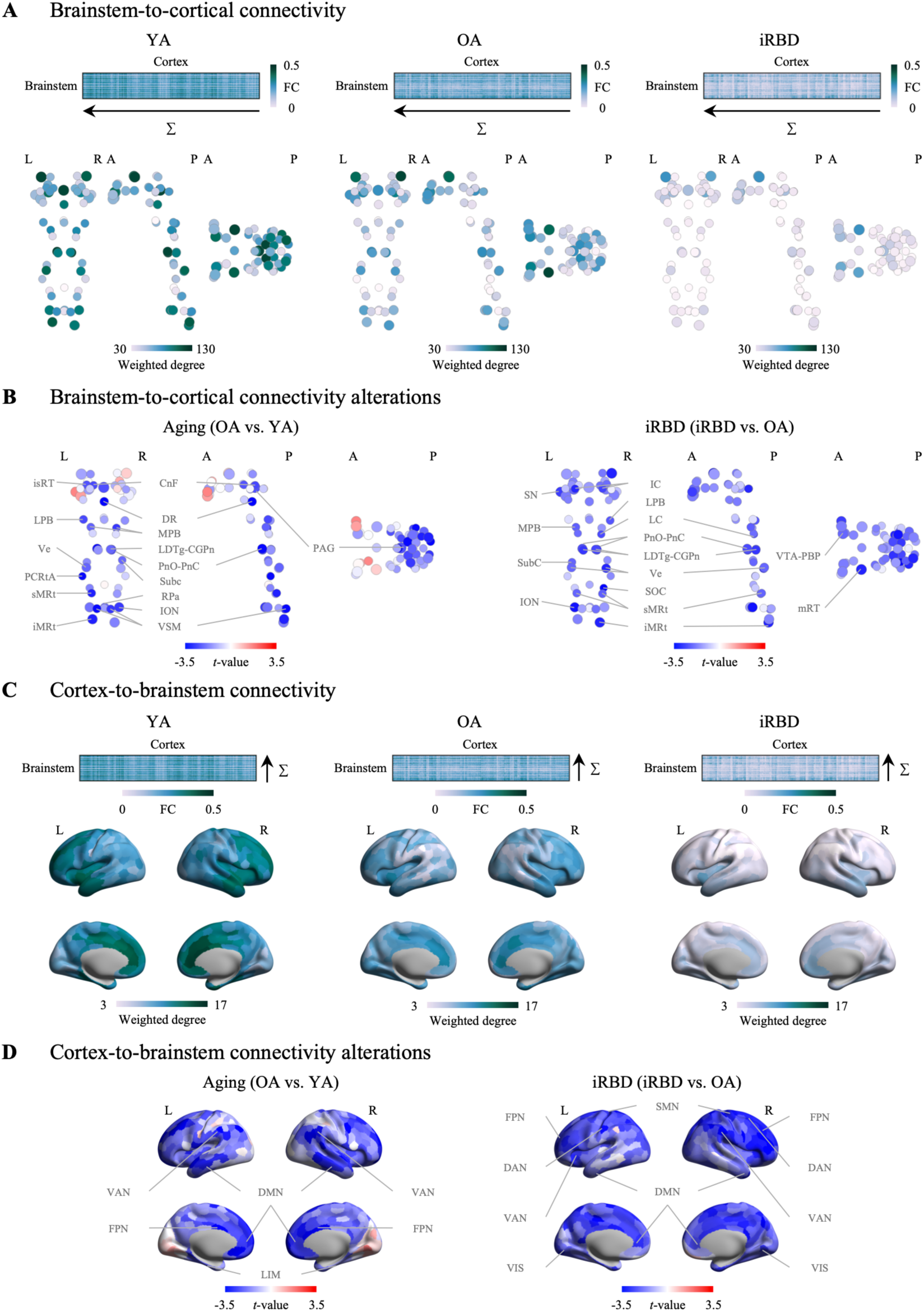
Brainstem-cortical weighted degree alterations in aging and iRBD. (A) Brainstem-to-cortical weighted degree was defined as the sum of each brainstem nucleus’s FC with all cortical regions for YA, OA and iRBD. Coronal, sagittal and horizontal views are shown with node size and color reflecting nucleus volume and weighted degree, respectively. (B) Group differences in brainstem-to-cortical weighted degree in aging (left) and iRBD (right). Node color presents the *t* values of weighted degree differences, and significant brainstem nuclei are labeled. (C) Cortex-to-brainstem weighted degree was calculated as the sum of each cortical region’s FC with all brainstem nuclei across groups. Color bars reflect FC strength (upper) and weighted degree (lower). (D) ROI-level cortex-to-brainstem weighted degree changes in aging (left) and iRBD (right). Color bars denote the *t* values of weighted degree changes, and significant functional networks are labeled.

**Figure 3.**
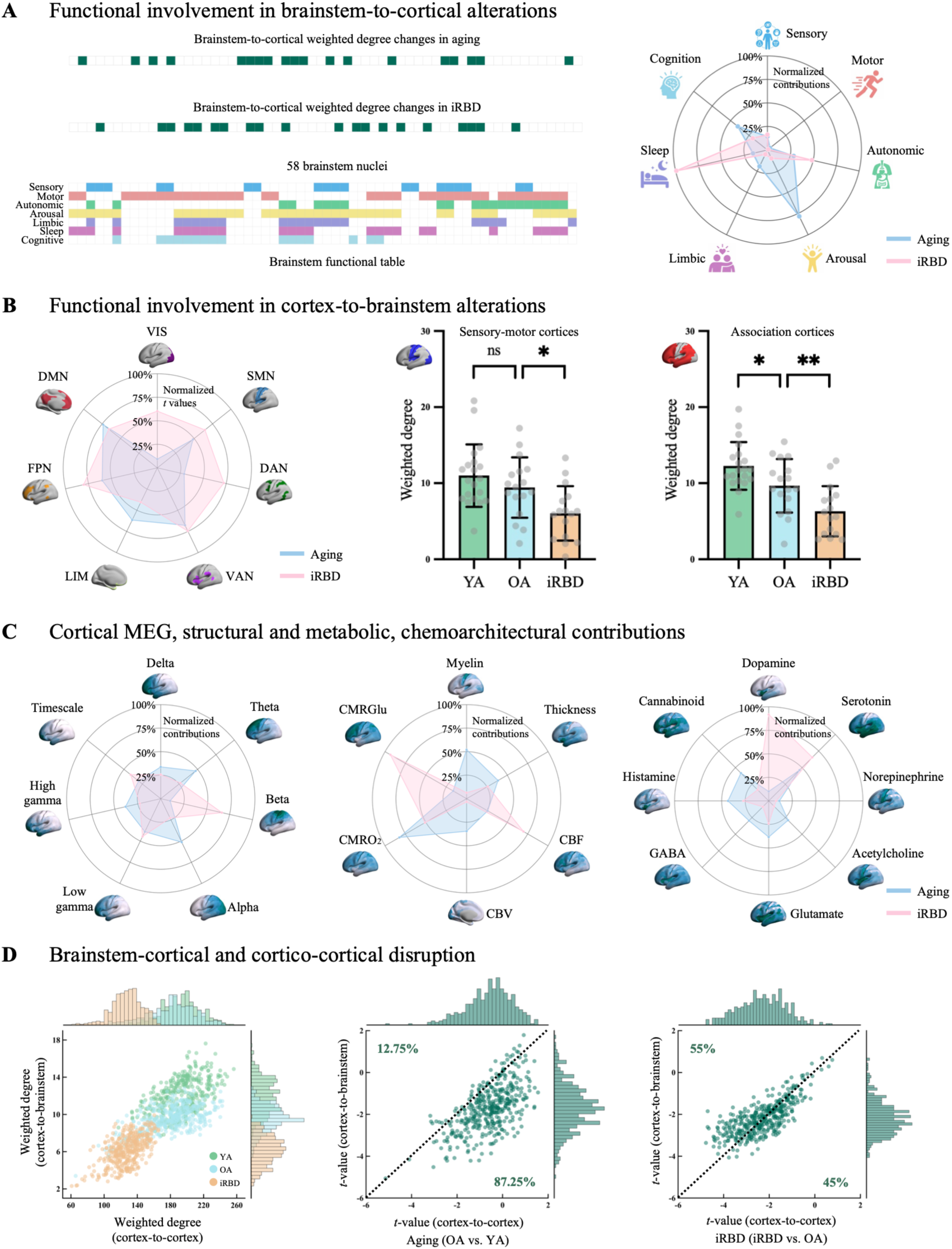
Neurobiological profiling of brainstem-to-cortical weighted degree alterations in aging and iRBD. (A) Binary maps of significant brainstem-to-cortical weighted degree changes in aging and iRBD (upper left) were used as dependent variables in logistic regression models. The seven columns of the binary matrix encoding brainstem functional domains (sensory, motor, autonomic, arousal, limbic, sleep and cognitive) were entered as independent variable, with colors denoting different functional domains (lower left) based on the available animal and human literatures (Supplementary Table S2). Percent contributions of each functional domain, quantified using dominance analysis, are summarized in the radar plot (right). Age- (*R^2^_adj_* = 0.36) and iRBD-related (*R^2^_adj_* = 0.49) contributions are shown in blue and pink, respectively. (B) Network-level cortex-to-brainstem weighted degree alterations across Yeo’s seven functional networks (left) and further integrated into sensory-motor and association cortices in aging and iRBD (right). (C) Separate multiple linear regression models were fitted between cortex-to-brainstem weighted degree alterations and cortical MEG dynamics, structure and metabolism, or neurotransmitter system density profiles for aging and iRBD. Dominance analysis was applied separately to the independent variables (cortical MEG dynamics, structural and metabolic features, or neurotransmitter receptors and transporters) to determine which cortical MEG dynamics (*R^2^_adj_* = 0.4 for aging and *R^2^_adj_* = 0.3 for iRBD; left), structural and metabolic features (*R^2^_adj_* = 0.44 for aging and *R^2^_adj_* = 0.19 for iRBD; middle), or neurotransmitter receptors/transporters (*R^2^_adj_* = 0.48 for aging and *R^2^_adj_* = 0.16 for iRBD; right) contributed most to the model fit in aging and iRBD. (D) Correspondence between cortex-to-cortex and cortex-to-brainstem weighted degree across groups (YA: *r* = 0.49; OA: *r* = 0.51; iRBD: *r* = 0.53, all *ps* < 0.001; left). Spatial correspondence between regional *t* values of cortex-to-cortex and cortex-to-brainstem weighted degree changes in aging (middle) and iRBD (right). Percentages indicate the proportions of cortical regions lying above and below the identity line, reflecting relatively greater cortex-to-cortex and cortex-to-brainstem weighted degree changes, respectively. Acronyms: VIS = visual network; SMN = somatomotor network; DAN = dorsal attention network; VAN = ventral attention network; LIM = limbic network; FPN = frontoparietal network; DMN = default mode network.

Cortex-to-brainstem weighted degree was then calculated by summing each cortical region’s FC across all brainstem nuclei (Figure 2C). Age-related reductions were predominantly localized to association cortex (FDR-corrected *p* = 0.02), whereas iRBD-related reductions involved both association cortex (FDR-corrected *p* = 0.009) and sensory-motor cortex (FDR-corrected *p* = 0.024; Figures 2D and 3B). The topography of cortical vulnerability affected by brainstem alterations showed distinct associations with multiple features of cortical organization. Cortical MEG dynamics showed stronger contributions of alpha power to age-related alterations (20.52%, *p_spin_* = 0.0361), whereas iRBD-related alterations were particularly aligned with beta power (26.61%, *p_spin_* = 0.031; Figure 3C). Structural features, including myelin and cortical thickness, contributed relatively modestly to both age- and iRBD-related alterations, whereas metabolic features contributed more prominently, with oxygen metabolism (32.97%, *p_spin_* = 0.011) contributing strongly to age-related alterations and cerebral blood flow (27.93%, *p_spin_* = 0.0498) and glucose metabolism (37.35%, *p_spin_* = 0.018) contributing strongly to iRBD-related alterations (Figure 3C). Neurotransmitter receptor and transporter distributions also revealed distinct patterns, with stronger contributions from dopamine (36.71%, *p_spin_* = 0.0242) and serotonin (26.19%, *p_spin_* = 0.0274) in iRBD and more uniform profiles in aging (Figure 3C). Finally, cortex-to-cortex and cortex-to-brainstem weighted degree showed strong spatial correspondence across groups (Figure 3D). Age-related changes were greater for cortex-to-cortex (87.25%) than cortex-to-brainstem (12.75%) weighted degree, whereas the opposite pattern emerged in iRBD, with greater cortex-to-brainstem (55%) than cortex-to-cortex alterations (45%; Figure 3D), consistent with preferential brainstem involvement in prodromal synucleinopathy^<u>7,29</u>^.

### Altered functional architecture of cortex-to-brainstem gradients

The principal cortex-to-brainstem gradient was derived from the similarity of cortical regions in their patterns of FC with the brainstem, after removal of the dominant brainstem-cortical connectivity pattern, extending from unimodal sensory-motor regions with negative scores to transmodal association regions with positive scores (Figure 4A). Across the whole cohort, brainstem-to-cortical weighted degree was calculated separately for negatively and positively scored cortical regions. Unimodal regions were most strongly connected with nuclei associated primarily with sensory (25.95%, *p_perm_* = 0.0354) and motor (25.02%, *p_perm_* = 0.0382) functions, including the inferior and superior colliculi and microcellular tegmental nucleus-parabigeminal nuclei in the midbrain, as well as the raphe pallidus and parvicellular reticular nucleus in the medulla. In contrast, transmodal regions were preferentially connected with functionally diverse nuclei involved in arousal (37.76%, *p_perm_* = 0.0164) function, including the periaqueductal gray, dorsal raphe, substantia nigra, mesencephalic reticular formation, caudal-rostral linear raphe, and ventral tegmental area-parabrachial pigmented nucleus complex in the midbrain, as well as the locus coeruleus, pedunculotegmental nucleus, laterodorsal tegmental nucleus-central gray of the rhombencephalon, and pontine reticular nucleus in the pons (Figure 4B).

**Figure 4.**
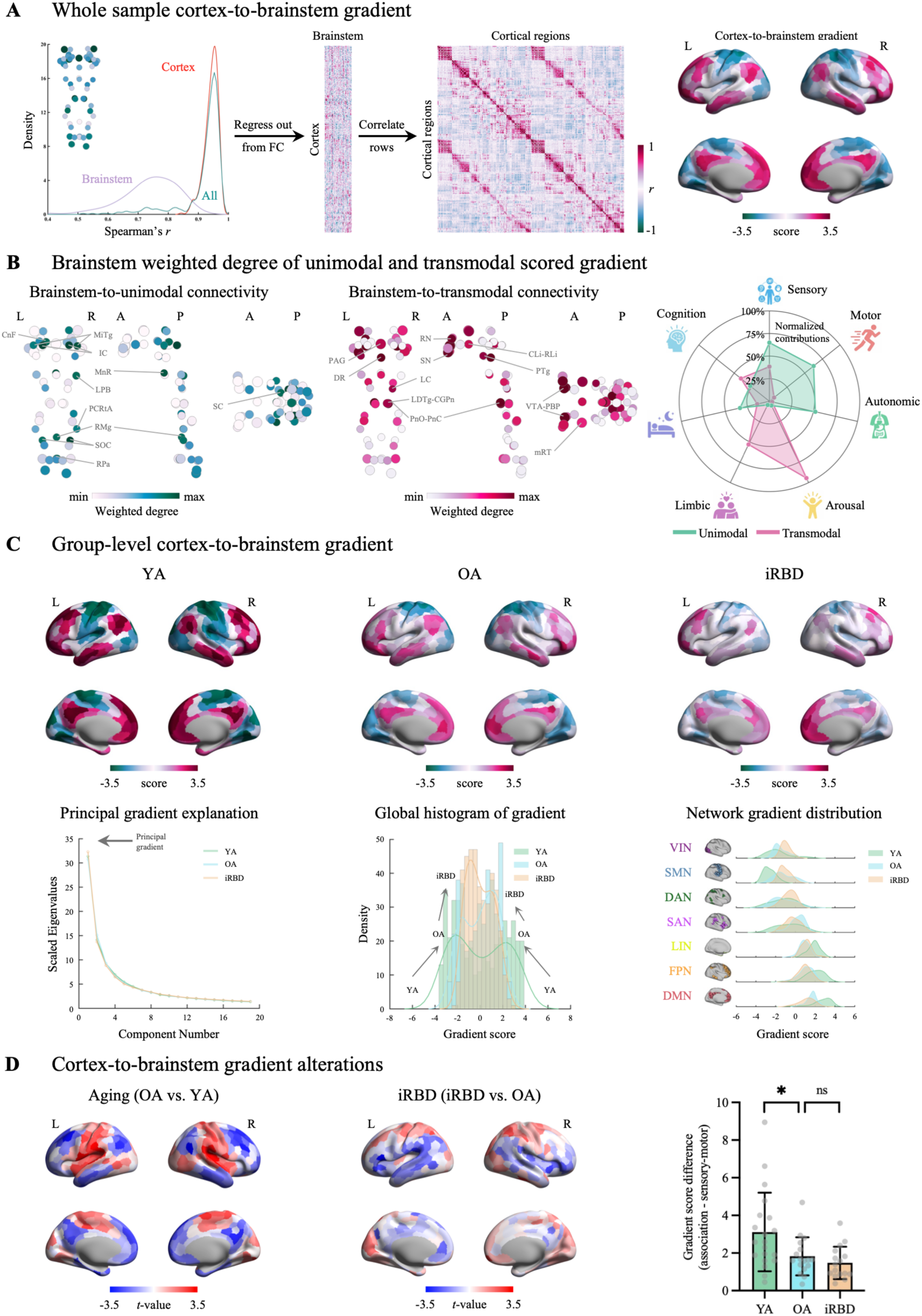
Altered hierarchical organization of cortex-to-brainstem functional connectivity in aging and iRBD. (A) For all 458 nodes (400 cortical and 58 brainstem), each node’s brainstem FC profile was correlated with the brainstem-to-cortical weighted degree to quantify the dominant shared connectivity pattern. The density distributions of Spearman’s *r* for brainstem (purple), cortical (red), and all nodes (blue; median *r* = 0.94; left). The brainstem map (weighted degree of brainstem-to-cortical FC) was regressed out of each cortical region’s brainstem FC pattern, resulting in a matrix (400 cortical regions × 58 brainstem nuclei) of FC residuals. The correlation matrix represents how similarly two cortical regions are functionally connected with the brainstem, above and beyond the dominant pattern of connectivity between the brainstem and cortex (middle). Diffusion map embedding was applied to the matrix, yielding the first gradient of cortex-to-brainstem FC (right). Color bars indicate the gradient score. (B) Brainstem weighted degree patterns were calculated as the sum of a brainstem nucleus’s FC with all negatively (left) or positively (middle) scored regions of the cortical gradient. Percent contributions of each functional domain, quantified using dominance analysis, are summarized in the radar plot (right). Brainstem-to-unimodal and brainstem-to-transmodal contributions are shown in dark green and pink, respectively. Coronal, sagittal and horizontal views are shown with node size and color reflecting nucleus volume and weighted degree, respectively. (C) Group-level cortex-to-brainstem gradients across groups (upper). The proportion of connectome-level variance explained by the top 19 diffusion embedding gradient components across groups (lower left). Global histogram of cortex-to-brainstem gradient scores across groups (lower middle). Functional network-based cortex-to-brainstem gradient scores across groups (lower right). (D) ROI-level *t*-statistical comparisons of cortex-to-brainstem gradient scores in aging (left) and iRBD (middle). Gradient score differences between association and sensory-motor cortex across groups (right). Color bars indicate the *t* values of gradient-score alterations. Error bars represent the SD. * *p* < 0.05, ** *p* < 0.01, *** *p* < 0.001, *ns* not significant.

Following alignment of subject-level gradients to the group-level template, gradient scores became progressively more concentrated around zero from YA to OA and iRBD. Both negatively scored unimodal regions and positively scored transmodal regions shifted toward the center of the gradient, reducing the distance between the two ends of the cortical distribution (Figure 4C). Accordingly, the separation between unimodal and transmodal regions differed significantly across groups (one-way ANOVA; *F*_(2,50)_ = 6.2, *p* = 0.004; Figure 4D). Group comparisons showed reduced unimodal-transmodal separation in aging (FDR-corrected *p* = 0.017), with no further significant difference in iRBD (FDR-corrected *p* = 0.501). Therefore, aging was associated with reduced spatial differentiation between unimodal and transmodal cortical regions in their patterns of connectivity with the brainstem, but not in iRBD.

### Spatial transcriptomic associations with brainstem-cortical alterations in aging and iRBD

Spatial transcriptomic patterns revealed that alterations in brainstem-to-cortical weighted degree were closely linked to regional gene expression in the brainstem. PLS1 explained 43.08% of the variance in age-related and 22.62% of the variance in iRBD-related weighted degree alterations. PLS1 scores showed significant spatial correspondence with regional *t*-statistic maps of weighted degree in both aging (*r* = 0.66, *p_perm_* = 0.0002; Figure 5A) and iRBD (*r* = 0.48, *p_perm_* = 0.0008; Figure 5B). Furthermore, GO enrichment indicated age-related weighted degree alterations were associated with biological processes involved in regulation of anatomical structure size and pattern specification process, whereas iRBD-related alterations were preferentially related to intermediate filament-based process, including intermediate filament cytoskeleton organization, intermediate filament bundle assembly, and neurofilament bundle assembly (FDR-corrected *ps* < 0.05; Figure 5C). KEGG pathway enrichment highlighted age-related associations with metabolic and mitochondrial pathways, as well as multiple neurodegenerative disease pathways, whereas iRBD-related alterations showed prominent enrichment in oxidative stress-related pathways, alongside convergent involvement of metabolic, mitochondrial, and neurodegenerative disease pathways (FDR-corrected *ps* < 0.05; Figure 5D), highlighting distinct and shared molecular pathways spatially associated with brainstem functional alterations across aging and iRBD.

**Figure 5.**
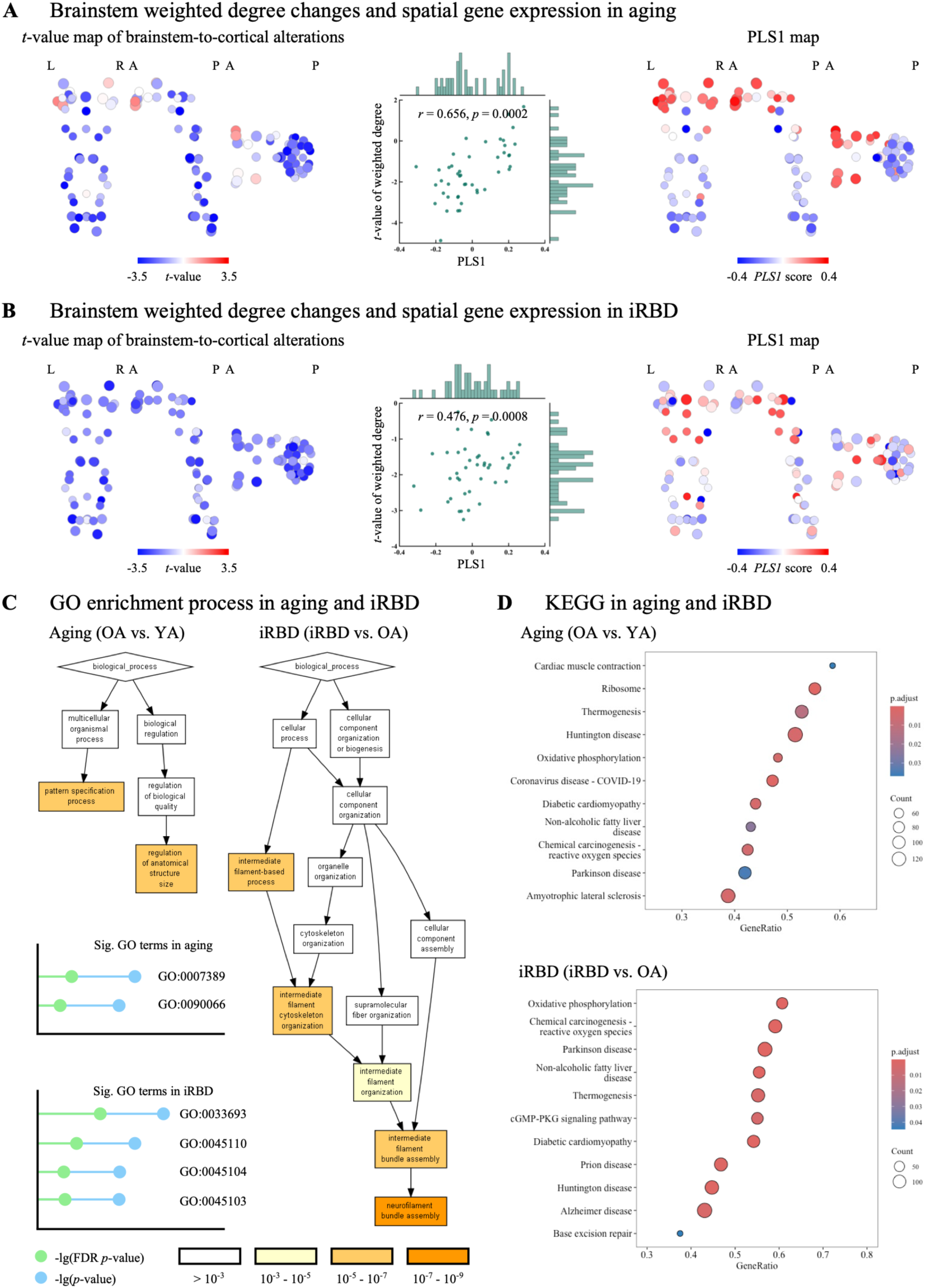
Spatial gene-expression patterns linked to brainstem weighted degree alterations. (A-B) Spatial correspondence (middle) between the *t*-map of brainstem-to-cortical weighted degree differences (left) and PLS1 scores of brainstem gene expression (right) in aging (A) and iRBD (B). (C) GO enrichment based on 15,633 genes ranked by PLS weights, shown separately for aging (left) and iRBD (right) and visualized as lollipop plots (green dots: FDR-corrected *p* < 0.05; blue dots: uncorrected *p* < 0.05). Age-related weighted degree alterations were associated with regulation of anatomical structure size (GO: 0090066; FDR-corrected *p* = 3.24×10^-2^) and pattern specification process in aging (GO: 0007389; FDR-corrected *p* = 5.69×10^-3^), whereas iRBD-related alterations were related to intermediate filament-based process in iRBD (GO: 0045103; FDR-corrected *p* = 1.63×10^-2^), including intermediate filament cytoskeleton organization (GO: 0045104; FDR-corrected *p* = 1.89×10^-2^), intermediate filament bundle assembly (GO: 0045110; FDR-corrected *p* = 2.84×10^-3^), and neurofilament bundle assembly (GO: 0033693; FDR-corrected *p* = 7.38×10^-5^). (D) KEGG pathway enrichment for aging (upper) and iRBD (lower). Color bars present the adjusted *p* value for pathway enrichment, and circle size indicates gene count contributing to each pathway.

### Robustness and sensitivity

The robustness of the main findings was further assessed using an alternative cortical parcellation (Schaefer 100-region atlas) and an independent 3-Tesla fMRI dataset from the PPMI. The Schaefer 100-region atlas reproduced the principal patterns observed with the primary parcellation, including regional FC, FC distributions and FC-distance relationships across groups, as well as age- and iRBD-related alterations in FC-distance relationships (Figure S5). Brainstem-to-cortical and cortex-to-brainstem weighted degree, together with their alterations in aging and iRBD, were similarly reproduced (Figure S6). In addition, FC patterns derived from the 7-Tesla dataset showed strong spatial correspondence with those from the independent 3-Tesla PPMI dataset. Group-averaged brainstem-brainstem FC was significantly correlated across datasets in OA (*r* = 0.31) and iRBD (*r* = 0.38), as were brainstem-cortical FC (OA: *r* = 0.32; iRBD: *r* = 0.47) and cortico-cortical FC (OA: *r* = 0.59; iRBD: *r* = 0.53, all *ps* < 0.001; Figure 6A). Group-averaged brainstem-to-cortical and cortex-to-brainstem connectivity also showed high cross-dataset correspondence (Figure 6B and 6C), supporting the generalizability across different scanners and processing pipelines.

**Figure 6.**
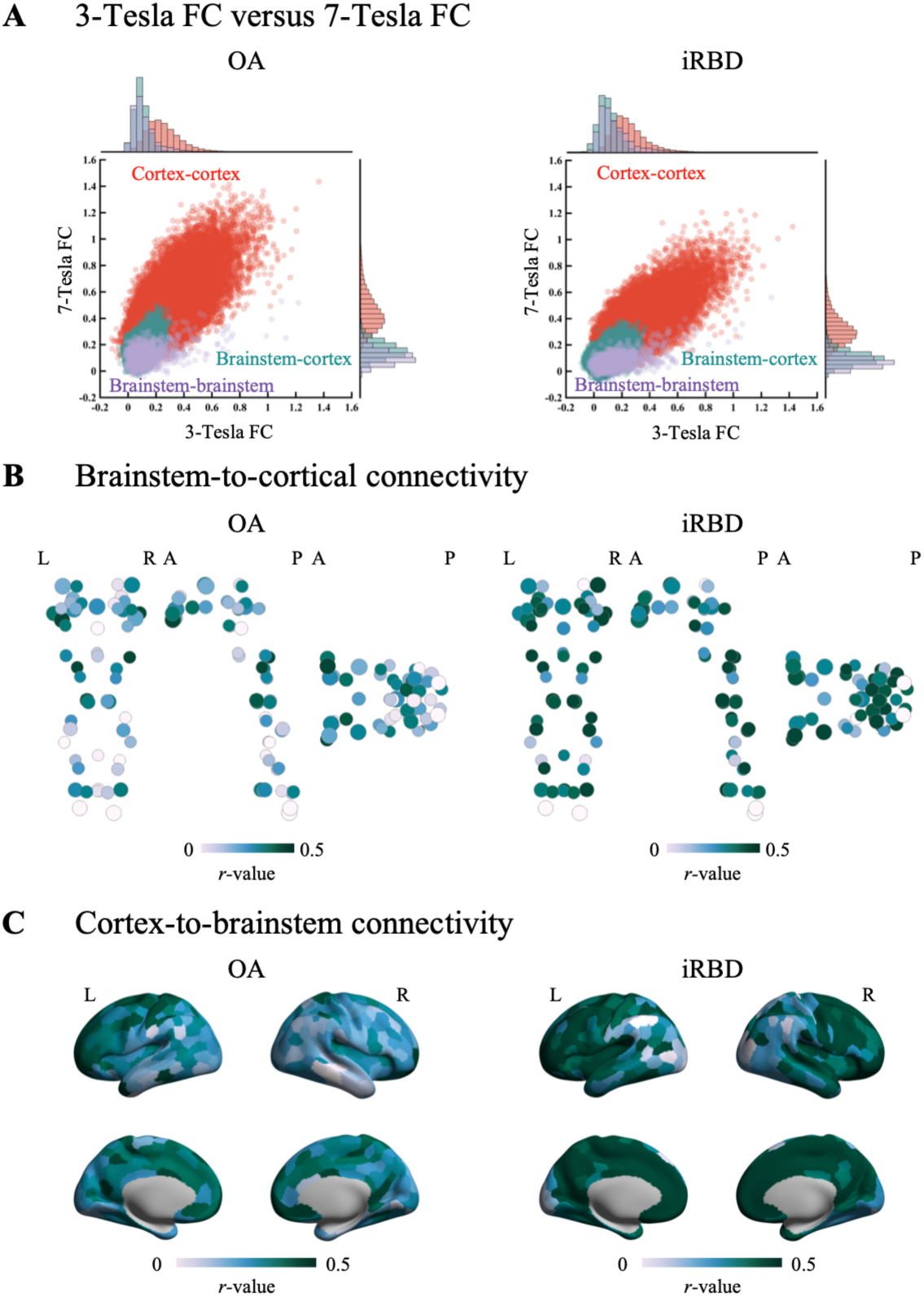
Robustness and translatability of 7-Tesla findings to 3-Tesla fMRI in the PPMI dataset. (A) Cross-dataset correspondence of regional FC in OA and iRBD. Scatter plots and marginal distributions show correlations between 7-Tesla and 3-Tesla data for brainstem-brainstem (purple), brainstem-cortical (green), and cortico-cortical (red) FC. (B) Spatial correspondence between 7-Tesla and 3-Tesla brainstem-to-cortical connectivity in OA and iRBD. Coronal, sagittal and horizontal views are shown, with node size indicating nucleus volume and node color indicating the correlation coefficient (*r*). (C) Spatial correspondence between 7-Tesla and 3-Tesla cortex-to-brainstem connectivity in OA and iRBD. Color bars indicate the correlation coefficient (*r*).

Finally, to assess individual variability, pairwise correlations of FC patterns were calculated across participants. FC patterns showed relatively low pairwise correlations, whereas brainstem-to-cortical and cortex-to-brainstem weighted degree was more conserved across individuals (Figure S7A), consistent with previous studies^<u>30</u>^ highlighting sensitivity of FC to individual differences. Importantly, individual variability quantified by standard deviation (σ) was stable across groups for brainstem-cortical FC, brainstem-to-cortical weighted degree, and cortex-to-brainstem weighted degree (Figure S7B-D).

## Discussion

In the present study, we mapped alterations in brainstem functional architecture across aging and prodromal synucleinopathy using ultra-high-field 7-Tesla resting-state fMRI in conjunction with a brainstem atlas of 58 nuclei. We identified that aging and iRBD shared preferential reductions of long-range brainstem-cortical connectivity but differed markedly in the neurobiological and molecular architectures associated with the affected pathways. Aging primarily involved brainstem nuclei supporting arousal and cognitive functions and their connections with higher-order association cortex, together with reduced differentiation between unimodal and transmodal cortical regions. In contrast, iRBD involved sleep- and autonomic-related nuclei and extended into lower-order sensory-motor cortex. Furthermore, we found that age-related cortical alterations in connectivity with the brainstem were more closely associated with alpha-band oscillations, oxygen metabolism, and more uniform chemoarchitectonic architecture, whereas iRBD-related alterations showed stronger associations with beta-band oscillations, cerebral blood flow, glucose metabolism, dopaminergic and serotonergic architectures. Regional gene-expression patterns were further associated with brainstem functional alterations, revealing partly shared metabolic, mitochondrial and neurodegenerative pathways, alongside distinct molecular annotations of age- and iRBD-related patterns. Together, current findings identify common vulnerability of long-range brainstem-cortical connectivity in aging and prodromal synucleinopathy, with connectivity affected through distinct biological processes.

First, we identified preferential vulnerability of long-range brainstem-cortical connectivity across aging and iRBD. Long-distance functional connections contribute to integration between spatially separated neural systems, while organization of the human connectome reflects a trade-off between efficient global information transfer and biological costs associated with maintaining long-distance anatomical connections^<u>31</u>^. Previous studies have reported reduced long-range FC and network efficiency, together with reduced differentiation between normally segregated networks^<u>32–35</u>^. Present findings extend distance-dependent vulnerability beyond cortico-cortical organization to functional interactions between the brainstem and cortex. Whereas brainstem-cortical FC showed relatively little dependence on anatomical distance in YA, increasing distance was associated with progressively weaker FC in OA and iRBD, indicating preferential disruption of spatially distant functional interactions. Long-distance interactions may be particularly important for the brainstem because brainstem nuclei participate in widely distributed ascending systems that regulate cortical activity and behavioral states. Maintaining long-distance communication requires substantial cellular and metabolic support, including mitochondrial energy production and axonal transport^<u>36</u>^. Age-related deterioration of mitochondrial function^<u>37,38</u>^ and α-synuclein-associated impairment of mitochondrial and axonal homeostasis in prodromal synucleinopathy^<u>39,40</u>^ may provide plausible biological explanations for the preferential vulnerability of long-range brainstem-cortical connectivity in aging and iRBD, respectively. Consistently, both age- and iRBD-related brainstem-cortical connectivity alterations were spatially enriched for mitochondrial and metabolic pathways.

Further, despite a shared vulnerability of long-range connectivity, aging and prodromal synucleinopathy reshaped brainstem-cortical organization through distinct systems-level patterns. Age-related vulnerability of brainstem hubs pointed to disruption of arousal and cognitive functions with higher-order cortical activity. In particular, alterations involved the dorsal raphe, which provides extensive serotonergic projections to forebrain regions, and the parabrachial complex, which promotes cortical arousal through ascending projections involving hypothalamic and basal-forebrain pathways, implicating major ascending regulatory circuits in aging^<u>21,41</u>^. Age-related alterations also involved the cuneiform nucleus, isthmic reticular formation, pontine reticular nuclei, and vestibular nuclei complex, which participate in interconnected arousal and motor systems, linking behavioral-state regulation with motor function^<u>42</u>^. Our findings are aligned with previous evidence of age-related weakening of ascending arousal connectivity with cortical association networks and its relationship with cognitive decline^<u>21</u>^. Interestingly, the locus coeruleus was not identified as an altered hub, potentially reflecting its previously reported non-linear lifespan trajectory^<u>43</u>^.

Likewise, we found that aging-related alterations in cortical FC with the brainstem preferentially involved association cortex. These cortical regions are predominantly situated toward the transmodal end of the principal cortical hierarchy, which extends from primary sensory-motor regions to higher-order integrative cortex^<u>44,45</u>^. Importantly, our findings further showed that brainstem connectivity is organized along the principal hierarchy, with transmodal cortical regions preferentially connected with arousal nuclei in the midbrain and pons, whereas unimodal cortical regions were more strongly connected with sensory and motor brainstem nuclei. The preferential involvement of association cortex in age-related brainstem-cortical alterations is supported by the established role of ascending neuromodulatory systems in regulating cortical activity and large-scale network integration^<u>46</u>^, together with human evidence that connectivity between ascending arousal systems and the default mode and salience networks declines with age^<u>21</u>^. Cortical organization is strongly differentiated along the sensory-association axis in healthy populations^<u>44,45</u>^, whereas aging is accompanied by reduced segregation among large-scale functional networks and differential reorganization of association relative to sensory-motor cortex^<u>33</u>^. Reduced unimodal-transmodal differentiation extends the concept of dedifferentiation beyond cortico-cortical organization by showing that cortical regions also become less distinct in their FC with the brainstem.

The biological architecture associated with cortex-to-brainstem connectivity alterations further reveals the multiscale cortical organization underlying age-related changes in brainstem connectivity. Alpha power contributed prominently, was consistent with evidence that alpha oscillatory activity undergoes marked and regionally heterogeneous changes across aging^<u>47,48</u>^, while oxygen metabolism align with the PET evidence of age-related changes in cerebral oxygen utilization and brain energy metabolism^<u>49,50</u>^. Neurotransmitter receptor and transporter distributions showed comparatively uniform contributions across systems, suggesting that age-related brainstem-cortical reorganization is embedded within distributed neurochemical architecture rather than preferentially aligned with a single neurotransmitter system. Collectively, aging appears to reshape brainstem-cortical communication beyond the loss of connectivity, altering how brainstem interactions are organized along the cortical functional hierarchy.

In contrast to aging, iRBD showed a distinct pattern of brainstem-cortical vulnerability, mainly involving nuclei associated with sleep and autonomic regulation. Alterations in the laterodorsal tegmental nucleus-central gray of the rhomboencephalon, subcoeruleus and pontine reticular formation implicated changes in interconnected pontine circuits supporting sleep-wake regulation and the generation and maintenance of REM sleep^<u>5,18–20</u>^. Consistent with previous 7-Tesla diffusion MRI evidence of altered connectivity between REM-on and medullary atonia-related regions in iRBD^<u>27</u>^, involvement of the inferior medullary reticular formation further supported alterations of pontomedullary circuits controlling REM sleep motor atonia, where pontine REM-active neurons engage inhibitory premotor neurons in the ventromedial medulla to suppress spinal motoneuron activity^<u>18–20,51</u>^. Beyond REM-control circuits, alterations of the substantia nigra, ventral tegmental area-parabrachial pigmented nucleus complex and locus coeruleus implicate dopaminergic and noradrenergic systems, aligning with neuromelanin-sensitive MRI, which provides complementary evidence of abnormalities in brainstem nuclei in individuals with iRBD^<u>52</u>^.

Cortical alterations associated with brainstem connectivity in iRBD extended beyond association cortex into sensory-motor cortex, distinguishing prodromal synucleinopathy from the predominantly association-cortical pattern observed with aging. Interestingly, cortico-cortical alterations were greater than cortex-to-brainstem alterations with aging, whereas the opposite pattern emerged in iRBD, supporting preferential brainstem involvement during prodromal synucleinopathy^<u>7,29</u>^. Extension into sensory-motor cortex may therefore reflect disruption of functional interactions between unimodal cortical regions and brainstem nuclei involved in sensory and motor processing, in line with fMRI studies identifying abnormalities across brainstem, basal ganglia and cortical sensory-motor regions in iRBD^<u>29,53</u>^. Notably, unimodal-transmodal differentiation was not further reduced in iRBD compared to healthy aging despite additional involvement of sensory-motor cortex. Prodromal synucleinopathy therefore primarily altered brainstem-cortical connectivity strength without further disrupting cortical hierarchical differentiation beyond age-related changes.

The greater contributions of cerebral blood flow and glucose metabolism in iRBD aligned with previous evidence of altered cortical perfusion and regional glucose metabolism^<u>54,55</u>^, suggesting that brainstem-cortical connectivity alterations preferentially involved cortical regions with distinct vascular and metabolic properties. Beta oscillations are strongly expressed across motor cortical and basal-ganglia circuits and become abnormally synchronized following dopaminergic dysfunction in parkinsonian states^<u>56</u>^. Spatial alignment between iRBD-related brainstem connectivity alterations and beta-band architecture may therefore relate disrupted brainstem-cortical interactions to neurophysiological processes supporting motor control. Neurotransmitter architecture further revealed stronger contributions from dopaminergic and serotonergic distributions, consistent with molecular imaging findings in iRBD and the predictive value of dopaminergic dysfunction for phenoconversion^<u>57–59</u>^. Gene-expression patterns spatially associated with iRBD-related brainstem alterations were enriched for intermediate-filament and neurofilament organization, linking affected brainstem regions to molecular architecture supporting axonal structure and transport. Neurofilaments are major neuronal intermediate filaments that maintain axonal caliber and structural integrity, while disruption of their homeostasis is associated with axonal dysfunction and neurodegeneration^<u>60,61</u>^. Collectively, iRBD appears to reshape brainstem-cortical communication through multisystem vulnerability extending into sensory-motor cortex, rather than simply amplifying age-related changes in cortical hierarchical organization.

Finally, the spatial correspondence between 7-Tesla connectivity patterns and independent 3-Tesla data supports the generalizability and cross-field-strength reproducibility of alterations in the functional architecture. This convergence highlights the complementary value of the two field strengths: 7-Tesla MRI enables more anatomically precise characterization of connectivity involving small brainstem nuclei, whereas 3-Tesla MRI provides a pathway for evaluating the resulting network-level features in larger and more heterogeneous clinical cohorts.

The important question raised by the present findings is why long-range brainstem-cortical pathways may be particularly vulnerable to aging and prodromal synucleinopathy. Consistent with the metabolic vulnerability of long-range axons^<u>37,38</u>^, the preferential reduction of long-range brainstem-cortical FC in the present study was accompanied by enrichment of metabolic and mitochondrial pathways in both aging and iRBD. Our findings extend previously described metabolic vulnerability of long-range axons to brainstem-cortical functional architecture and suggest that the high biological demands required to maintain spatially distributed communication may contribute to its preferential vulnerability^<u>31,37,38</u>^. However, shared long-range vulnerability did not lead to equivalent network reorganization across aging and iRBD, suggesting that prodromal synucleinopathy does not simply amplify age-related hierarchical dedifferentiation but instead involves additional functional systems within the altered aging architecture. Such divergence is consistent with evidence that neurodegenerative progression depends not only on network connectivity but also on regional biological vulnerability^<u>62</u>^. Our multiscale findings provide complementary evidence for this principle in brainstem functional organization, with aging and iRBD affecting shared long-range architecture while the affected pathways were associated with distinct neurobiological and molecular profiles. The distinction may also help explain how iRBD, despite being clinically defined by abnormal REM sleep behavior, precedes neurodegenerative syndromes involving motor, autonomic and cognitive dysfunction^<u>5,16,17</u>^. Longitudinal studies demonstrate that multiple motor and non-motor abnormalities evolve during the iRBD stage before phenoconversion^<u>16,17</u>^, while neuroimaging studies identify structural and functional abnormalities extending beyond canonical REM-control circuits^<u>27,29,53</u>^. Our findings similarly identify alterations beyond REM-related brainstem circuits. Clinical onset may therefore not coincide with the extent of network dysfunction, as a relatively specific sleep phenotype may become clinically apparent while functional alterations are already distributed across systems relevant to later motor and non-motor manifestations^<u>16,17,29</u>^. This broader involvement may be clinically significant, as the substantia nigra, locus coeruleus and medullary reticular formation are prominently implicated in Parkinson’s disease^<u>63</u>^, dementia with Lewy bodies^<u>64</u>^ and multiple system atrophy^<u>65</u>^, raising the possibility that their involvement during the iRBD stage may relate to emerging motor, autonomic and cognitive abnormalities and, ultimately, to distinct phenoconversion trajectories. Longitudinal studies comparing iRBD with manifest synucleinopathies will be important to determine whether these brainstem systems differentiate distinct phenoconversion trajectories.

Several methodological limitations need to be noted regarding the present study. First, precise neuroimaging of the human brainstem nuclei is inherently challenging due to their deep anatomical location, proximity to major vasculature and CSF, irregular morphology, and small nuclear size. Although the present data underwent brainstem-optimized physiological noise correction and the delineation of brainstem nuclei was carefully validated, brainstem fMRI methodology is still evolving, and best practices continue to be refined. Second, the temporal resolution of the 7-Tesla fMRI was limited to 2.5s. The trade-off was necessary to accommodate the number of slices and high spatial resolution required to reliably resolve small brainstem nuclei. Third, FC measures derived from resting-state fMRI reflect indirect and potentially polysynaptic interactions rather than direct anatomical connections^<u>66</u>^. Therefore, the brainstem-cortical connectivity patterns should not be interpreted as evidence of monosynaptic pathways. Fourth, the present study primarily focused on group-level alterations of brainstem-cortical connectivity in aging and iRBD, rather than on inter-individual variability. Notably, the magnitude of individual variability was stable across groups. As acquisition protocols and processing pipelines for brainstem fMRI become increasingly standardized, future studies with larger samples will enable investigation of whether individual differences in brainstem-cortical connectivity predict disease-related phenotypes and inform personalized biomarkers and intervention strategies.

In summary, we delineated alterations in long-range brainstem-cortical functional architecture in aging and iRBD. Our findings provide multiscale evidence linking nucleus-level brainstem vulnerability to functional, neurobiological and molecular organization in aging and prodromal synucleinopathy, highlighting the potential of brainstem pathways as early biomarkers and providing a framework for understanding the biological mechanisms underlying large-scale network dysfunction in aging and prodromal synucleinopathy.

## Methods

### Participants and study design

Twenty young adults (YA; age: 29.35 ± 5.4 years; 10 females), seventeen older adults (OA; age: 64.41 ± 4.15 years; 3 females), and sixteen individuals with iRBD (age: 67.19 ± 5.96 years; 3 females) were recruited and underwent 7-Tesla MRI. Sex distribution did not differ significantly among groups (Fisher’s exact test, *p* = 0.067). As expected, YA were significantly younger than both OA and iRBD participants (FDR-corrected *ps* < 0.001), whereas no significant age difference was found between OA and iRBD (FDR-corrected *p* = 0.133). All participants provided written informed consent in accordance with protocols approved by the Institutional Review Board of Massachusetts General Hospital. Inclusion criteria were consistent across groups and required: no history of neurological trauma, normal or corrected-to-normal vision, absence of claustrophobia, capacity to perform experimental tasks, no metal implants or prior exposure to metal fragments, ability to provide informed consent, and a head circumference under 61 cm (to accommodate the 7-Tesla MRI coil). Exclusion criteria for YA and OA included any history of sleep or neurological disorders. Individuals with iRBD were evaluated by a board-certified neurologist (A.V.). Diagnosis followed the International Classification of Sleep Disorders (ICSD-3), based on repeated episodes of sleep-related vocalizations and/or complex motor behaviors during sleep, and the exclusion of alternative causes (e.g., epilepsy). None of the individuals with iRBD showed clinical signs of parkinsonism, dementia, or other neurological comorbidities upon examination (Supplementary Table S1).

During scanning, participants were instructed to lie still with their eyes closed while remaining awake. To minimize head motion, foam padding was placed around the head. Cardiac and respiratory cycles were simultaneously recorded using a piezoelectric finger pulse sensor (ADInstruments, Colorado Springs, CO, USA) and a piezoelectric respiratory bellow (UFI, Morro Bay, CA, USA) positioned around the chest, respectively. Physiological signals were digitized at 1000 Hz using a PowerLab data acquisition system (PowerLab 4/SP, ADInstruments, New Zealand) and LabChart software (LabChart, ADInstruments, New Zealand). The same acquisition system also recorded the MR scanner trigger pulses synchronized with the acquisition of each image volume. Wakefulness was assessed using responses to questions delivered via an MRI-compatible buzzer system immediately before and after each run.

### 7-Tesla fMRI data acquisition

Resting-state fMRI data were acquired using a 7-Tesla MRI scanner (Magnetom, Siemens Healthineers, Erlangen, Germany) with a custom-built 32-channel receive coil and volume transmit coil^<u>67</u>^. Brainstem-specific custom protocols were developed and used for the 7-Tesla MRI acquisition and subsequent processing^<u>42,68</u>^. For each participant, functional gradient-echo echo-planar images (EPIs) were acquired with the following parameters: isotropic voxel size = 1.1 mm, matrix size = 180×240, GRAPPA factor = 3, nominal echo-spacing = 0.82 ms, bandwidth = 1488 Hz/Px, number of slices = 123, slice orientation = sagittal, slice-acquisition order = interleaved, echo time (TE) = 32 ms, repetition time (TR) = 2.5 s, flip angle (FA) = 75°, simultaneous-multi-slice factor = 3, number of repetitions = 210, phase-encoding direction = anterior-posterior, and acquisition-time = 9 min 25 s. To correct for geometric distortion of the functional images, field-map images were collected with the following parameters: isotropic voxel size = 2.0 mm, matrix size = 116×132, bandwidth = 1515 Hz/Px, number of slices = 80, slice orientation = sagittal, slice-acquisition order = interleaved, TEs = [3.00, 4.02] ms, TR = 570 ms, FA = 36°, simultaneous-multi-slice factor = 3, phase-encoding direction = anterior-posterior, acquisition time = 2 min 15 s. Finally, high-resolution anatomical T_1_-weighted multi-echo MPRAGE (MEMPRAGE) images were acquired to enable precise parcellation of brainstem, subcortical, and cortical regions and accurate coregistration of fMRI data to stereotactic space, using the following parameters: isotropic voxel size = 0.75 mm, TR = 2.53 s, TEs = [2.39, 5.62] ms, inversion-time = 1.1 s, FA = 7°, field of view = 240×240×240 mm^3^, bandwidth = 332 Hz/Px, GRAPPA factor = 2, slice orientation = sagittal, slice-acquisition order = anterior-posterior, acquisition time = 6 min 34 s.

### 7-Tesla fMRI data pre-processing

Resting-state fMRI data were pre-processed using a brainstem-optimized pipeline tailored for 7-Tesla MRI^<u>42,68</u>^. For anatomical preprocessing, the root mean square across echo times was computed from each MEMPRAGE image, which was then reoriented to standard right-posterior-inferior (RPI) orientation. Bias-field correction was applied using SPM12 toolbox (https://www.fil.ion.ucl.ac.uk/spm/software/spm12/), followed by skull stripping and image cropping using FSL software (version 5.0.7; https://fsl.fmrib.ox.ac.uk/fsl/). Anatomical images were spatially normalized to the MNI152NLin6Asym 1-mm isotropic template space using Advanced Normalization Tools (ANTs, version 2.2.0; https://github.com/ANTsX/ANTs), incorporating affine and nonlinear transformations optimized for accurate alignment of brainstem anatomy (detailed anatomical coregistration procedures are provided in the Supplementary Material; Figure S1A and S1B).

Field-map preprocessing using a hybrid approach was performed using SPM12 toolbox and FSL software. Specifically, magnitude images were rotated to RPI orientation, bias-field corrected, and skull-stripped. Brain masks were manually edited to ensure complete inclusion of brainstem structures, particularly the lower medulla, and to exclude residual noisy non-brain voxels. Phase images were subsequently rotated to RPI orientation, converted to angular frequency (radians/sec), masked using the corrected brain masks, and smoothed with a 3D Gaussian kernel (σ = 4 mm). Magnitude images were warped to match the angular frequency images (FSL *fugue* command) and used to linearly register the angular frequency image to the fMRI (FSL *flirt* command). Finally, angular frequency images registered to the fMRI were computed and generated individualized warps for fMRI distortion correction (FSL *fugue* command).

Physiological noise was corrected using RETROICOR^<u>69</u>^, implemented in custom-built MATLAB (R2017a, MathWorks, Natick, MA, USA) functions adapted to the slice acquisition order (Figure S1C). Functional images were slice-time corrected by temporal interpolation and reoriented to RPI orientation. Motion correction was performed relative to the first volume using FreeSurfer software (version 7.4.1; https://surfer.nmr.mgh.harvard.edu/fswiki). Six time-series describing rigid motion (three translations and three rotations) of the head were saved for subsequent use as nuisance regressors. To obtain a reference fMRI volume for the coregistration to the MPRAGE image with enhanced spatial signal-to-noise ratio, the first run of the motion-corrected fMRI data was time-averaged, bias-field corrected, and corrected for field-map-based geometric distortions. Then, the resulting reference fMRI was coregistered to the MEMPRAGE using AFNI (version 19.3.16; https://afni.nimh.nih.gov/) with a two-stage procedure consisting of affine alignment (AFNI *3dAllineate* command), followed by nonlinear registration (AFNI *3dQwarp* command) with multi-resolution refinement and edge-based weighting optimized for brainstem structures. To minimize repeated spatial interpolation and preserve spatial resolution, the affine and nonlinear fMRI-to-MEMPRAGE transformations were concatenated with the motion and distortion-correction transformations and applied to the fMRI images in a single interpolation step (AFNI *3dNwarpApply* command). Next, nuisance regressors were computed and regressed from the fMRI time series to reduce motion- and physiology-related signal fluctuations, with particular attention to the brainstem. Regressors included six rigid-body motion parameters, a respiratory volume per time (RVT) regressor convolved with a respiratory response function^<u>70</u>^, a heart rate (HR) regressor convolved with a cardiac response function^<u>71</u>^, and five components derived from principal component analysis on voxels within a cerebrospinal fluid (CSF) mask adjacent to the brainstem. The CSF mask was defined by thresholding the temporal standard deviation of the EPI signal within a 5.3 × 6.5 × 7.5 cm^3^ cuboid surrounding the brainstem and encompassed the lower portion of the third ventricle, the cerebral aqueduct, and the fourth ventricle. Previous studies have shown that the removal of CSF-related signal fluctuations, particularly those arising from the cerebral aqueduct and fourth ventricle, is important for minimizing physiological contamination in FC analysis of adjacent brainstem nuclei^<u>72,73</u>^. Finally, fMRI time series were normalized to percent signal change by dividing each voxel time series by its temporal mean and multiplying by 100, then were temporal band-pass filtered between 0.01 Hz and 0.1 Hz (AFNI *3dBandpass* command), and the residual mean signal was removed.

### Brainstem nuclei segmentation

Brainstem nuclei were delineated using the Brainstem Navigator (version 1.1; https://www.nitrc.org/projects/brainstemnavig), a probabilistic atlas of the *in-vivo* human brainstem recently developed by our team. The atlas comprises 58 brainstem labels (8 midline and 25 bilateral nuclei), and 8 diencephalic labels (4 bilateral diencephalic nuclei). All nuclei are represented in MNI152NLin6Asym 1-mm^3^ space (matrix size: 182 × 218 × 182). Detailed methods for atlas construction and validation have been reported previously^<u>27,74–77</u>^.

In brief, the atlas was constructed from ultra-high-field 7-Tesla MRI data acquired from 12 healthy adults (age: 28 ± 1 years; 6 females), including both T_2_-weighted and diffusion-weighted images. Fractional anisotropy (FA) maps derived from diffusion-weighted images, together with high-contrast T_2_-weighted images, were used to provide complementary contrasts for the delineation of brainstem nuclei boundaries. Initial segmentations targeted the raphe nuclei (median, dorsal, and magnus), periaqueductal gray, substantia nigra, and red nuclei^<u>74</u>^. A semi-automated clustering approach was applied at the individual-participant level to either FA or T_2_-weighted images, depending on the brainstem structure of interest. Manual refinement was performed when clusters encompassed multiple nuclei, guided by established anatomical landmarks. Individual binary masks for each nucleus were transformed to MNI152NLin6Asym 1-mm isotropic template space and averaged to generate group-level probabilistic maps, with each voxel value representing the proportion of participants in whom the voxel was assigned to the corresponding nucleus; thus, a value of 1 indicated agreement across all participants. Although single-subject segmentations were non-overlapping, minimal spatial overlap may appear in group-averaged templates depending on the threshold applied. Subsequent work expanded the atlas across the brainstem and diencephalon. The mesopontine tegmental nuclei (cuneiform nucleus, pedunculotegmental nucleus, oral pontine reticular nucleus, paramedian raphe, and caudal-linear raphe) were delineated using the same pipeline^<u>75</u>^. These segmentations were further validated through expert manual annotations by a neurosurgeon referencing multimodal image contrasts and anatomical landmarks, as well as cross-validation with a postmortem histologically defined atlas^<u>3</u>^. For subsequent nuclei, expert manual segmentation was conducted independently by multiple raters, with final labels defined by the intersection of rater-defined masks. Probabilistic maps were then generated by averaging the segmentations across individuals, and all structures were verified against the Paxinos atlas^<u>3</u>^. Further extensions included delineation of the inferior and superior colliculi and the superior olivary complex, alongside diencephalic thalamic nuclei such as the medial and lateral geniculate nuclei^<u>76</u>^. Singh et al.^<u>78</u>^ defined the medial and lateral parabrachial nuclei, vestibular nuclei complex, and the medullary viscero-sensory-motor nuclei, with additional histological validation of the parabrachial nuclei. Furthermore, Singh et al.^<u>77</u>^ subsequently delineated the mesencephalic and isthmic reticular formations, microcellular tegmental nucleus, ventral tegmental area, and the caudal-rostral linear raphe nucleus complex. Finally, García-Gomar et al.^<u>27</u>^ contributed segmentations of the raphe obscurus, raphe pallidus, locus coeruleus, subcoeruleus, laterodorsal tegmental nucleus-central gray of the rhombencephalon, inferior and superior medullary reticular formation, and the pontine reticular nucleus (oral and caudal part).

### Functional network construction

Individual-level functional connectomes were constructed using the 400-region Schaefer cortical atlas^<u>28</u>^ in combination with the 58-nuclei labels of the Brainstem Navigator atlas (Figure 1A and Figure S2). Spatial alignment of the fMRI data and atlas labels to MNI space was carefully inspected across multiple slices encompassing the brainstem for each participant in each group. Minor distortions in susceptibility-prone areas, such as the anterior surface of the pons, had no impact on FC construction, as no nodes were in the affected anterior pontine regions (Figure S3). For each participant, regional mean time series were extracted from the unsmoothed functional images registered to MNI space. Pairwise Pearson’s correlation coefficients were computed between the mean time series of all pairs of regions, resulting in a 458 × 458 FC matrix. FC values were subsequently Fisher z-transformed to improve the normality of their sampling distribution. To evaluate the spatial constraints shaping FC^<u>79</u>^, Spearman’s correlation coefficients between FC strength and anatomical (Euclidean) distance were computed separately for each group. The distribution of brainstem-cortical connectivity was characterized using weighted degree computed from the 58 x 400 brainstem-cortical FC matrix by summing FC values across cortical regions for each brainstem nucleus (“brainstem-to-cortical” weighted degree) and across brainstem nuclei for each cortical region (“cortex-to-brainstem” weighted degree)^<u>80</u>^. These measures therefore quantify connectivity strength from the perspective of brainstem and cortical nodes, respectively, rather than directional functional interactions. Cortical findings were aggregated within Yeo’s seven functional networks^<u>81</u>^ and projected onto the cortical surface using Connectome Workbench (version 2.1.0; https://www.humanconnectome.org/software/get-connectome-workbench). Network-level findings were averaged within Yeo’s seven functional cortical networks and classified into lower-order sensory-motor (visual and somatomotor networks) and higher-order association cortical networks (dorsal attention, ventral attention, limbic, frontoparietal and default mode networks) following prior definitions^<u>30,45</u>^. Age-related alterations were assessed by comparing OA with YA, whereas iRBD-related changes were evaluated by comparing iRBD with OA.

To assess the stability and generalizability of the findings, inter-individual variability in FC and in brainstem-to-cortical and cortex-to-brainstem weighted degrees were quantified using the across-participant standard deviation (SD; σ). The main findings were further evaluated using the 100-region Schaefer cortical atlas to assess their robustness to cortical parcellation resolution. In addition, an expanded parcellation was evaluated that incorporated subcortical regions, including 4 bilateral diencephalic nuclei from the Brainstem Navigator, the hypothalamus^<u>82</u>^, and the ventral and dorsal spinal cord^<u>27</u>^, together with 8 bilateral FreeSurfer-derived subcortical structures^<u>83</u>^.

### Functional and neurobiological characterization of altered brainstem-cortical connectivity

To identify the functional and neurobiological features underlying brainstem-cortical connectivity alterations in aging and iRBD, we assessed two complementary components of the altered connectivity architecture: (i) which functional domains characterized brainstem nuclei showing altered connectivity with the cortex, and (ii) which neurobiological properties characterized cortical regions showing altered connectivity with the brainstem. For brainstem-to-cortical weighted degree, seven functional domains were defined based on a systematic literature review, comprising sensory, motor, autonomic, arousal, limbic, sleep, and cognitive functions. A binary nucleus-function matrix was constructed across the 58 brainstem nuclei, with a value of 1 indicating literature-based evidence for involvement of a nucleus in a given functional domain and 0 indicating no reported evidence (Supplementary Table S2). Brainstem nuclei showing statistically significant alterations in brainstem-to-cortical weighted degree were coded as 1 and nuclei without significant alterations as 0, generating separate binary response variables for aging and iRBD. Binomial logistic regression was used to estimate the probability of a significant brainstem-to-cortical weighted degree alteration as a function of the seven functional domains. Relative predictor importance was quantified using dominance analysis for the logistic regression^<u>84</u>^, with McFadden’s *R*²^<u>85</u>^ as the index of model fit. For *p* predictors, all 2*^p^* - 1 possible non-empty predictor subsets were evaluated. The contribution of each functional domain was defined as the increase in McFadden’s *R*² produced by adding the predictor to a given subset model. Contributions were calculated across all possible subsets of the remaining predictors, averaged within each model size and subsequently across model sizes to derive the general dominance measure. General dominance measures sum to the McFadden’s *R*² of the full model, thereby partitioning overall model fit among the seven functional domains. Each general dominance measure was normalized by the full-model McFadden’s *R*² and expressed as a percentage contribution, representing the relative importance of each functional domain in explaining brainstem-to-cortical weighted degree alterations in aging and iRBD.

For cortex-to-brainstem weighted degree, normative brain maps capturing multiple scales of cortical organization were obtained from the neuromaps toolbox (version 0.0.7; https://github.com/netneurolab/neuromaps)^<u>86</u>^. Cortical annotations encompassed the following neurobiological domains: (i) seven neurophysiological maps (canonical magnetoencephalography (MEG) frequency-band power in the delta, theta, alpha, beta, low-gamma and high-gamma frequency bands, together with intrinsic neural timescale); (ii) two structural maps (cortical thickness and the T_1_-weighted/T_2_-weighted MRI ratio as a proxy for intracortical myelin), and four metabolic maps (cerebral blood flow, cerebral blood volume, cerebral metabolic rate of oxygen and glucose metabolism); and (iii) 18 PET-derived neurotransmitter receptor and transporter density maps spanning dopamine (D_2_ and DAT), norepinephrine (NET), serotonin (5-HT_1A_, 5-HT_1B_, 5-HT_2A_, 5-HT_4_, 5-HT_6_, 5-HTT), acetylcholine (α_4_β_2_, M1, VAChT), glutamate (mGluR_5_ and NMDA), GABA (GABA_A_), histamine (H_3_), and cannabinoid (CB_1_) systems. All annotations were parcellated according to the 400-region Schaefer atlas. To account for the distinct biological scales represented by these annotations and to avoid combining heterogeneous cortical features within a single model, dominance analyses were performed separately for the three feature sets (i)-(iii). Each feature set was analyzed separately for aging- and iRBD-related cortex-to-brainstem weighted degree alterations. Multiple linear regression models combined with dominance analysis^<u>87</u>^ were applied within each feature set to estimate the relative contribution of each cortical feature in explaining the spatial pattern of cortex-to-brainstem weighted degree alterations. For *p* predictors within a given feature set, all 2*^p^* - 1 possible non-empty predictor subsets were evaluated. Predictor-specific contribution was defined as the increase in *R*² produced by adding the predictor to every subset model not already containing it. Total dominance was calculated as the average incremental contribution across all relevant subset models, partitioning the overall model fit among predictors while accounting for shared variance among cortical features. Interactional dominance was defined as the increase in *R*² produced by adding a predictor to the model containing all remaining predictors within the same feature set, representing its unique incremental contribution in the presence of all other cortical features. Total and interactional dominance were normalized by the adjusted *R*² of the full model, with normalized total dominance expressed as the percentage contribution of each cortical feature in explaining the spatial pattern of cortex-to-brainstem weighted degree alterations.

### Cortex-to-brainstem connectivity gradients

Cortex-to-brainstem connectivity gradients were derived to capture hierarchical variation in cortical patterns of connectivity with the brainstem, beyond the dominant connectivity patterns. For each participant, global connectivity strength was first removed by regressing out the mean brainstem-to-cortical weighted degree from the brainstem-cortical FC matrix, thereby removing variance related to overall connectivity strength while preserving finer-grained distinctions in FC organization. Following regression, pairwise cosine similarity was calculated across the cortical columns of brainstem-cortical FC profiles, resulting in a cortex × cortex similarity matrix. Diffusion map embedding, a nonlinear dimensionality reduction approach that identifies principal gradients of functional organization^<u>88</u>^, was then applied to each participant’s similarity matrix. Specifically, diffusion embedding was performed by computing squared Euclidean distances from min-max normalized input matrices, followed by application of a Gaussian kernel (σ = 1), α-normalization (α = 1), symmetric normalization, and singular value decomposition. Gradient eigenvectors were sign-aligned to ensure directional consistency across participants, and the first non-trivial component was retained as the principal gradient of cortex-to-brainstem functional architecture. This gradient captures the dominant continuous axis along which cortical regions differ in their relative patterns of connectivity with brainstem nuclei. Gradient scores were subsequently averaged within Yeo’s seven functional networks and further grouped into lower-order sensory-motor and higher-order association cortical networks to evaluate group differences.

### Spatial transcriptomic mapping of altered brainstem-cortical connectivity

To investigate molecular signatures associated with brainstem alterations in aging and iRBD, spatial patterns of altered brainstem-to-cortical weighted degree were related to regional gene expression profiles. Gene expression data were obtained from the Allen Human Brain Atlas (AHBA; https://human.brain-map.org/), which comprises microarray expression data from six neurotypical adult donors, and were subsequently mapped to brainstem regions defined by the Brainstem Navigator using the abagen toolkit (https://github.com/rmarkello/abagen). Following preprocessing and regional mapping, expression data for 15,633 genes across brainstem nuclei were obtained. Partial least squares (PLS) regression was used to identify multivariate gene expression patterns associated with the spatial distribution of brainstem-to-cortical weighted degree alterations in aging and iRBD. Specifically, separate PLS analyses were performed for aging and iRBD, with regional *t*-statistic maps from the corresponding group comparisons in brainstem-to-cortical weighted degree serving as dependent variables, and regional gene expression profiles serving as independent variables. Given the high dimensionality of the gene expression data relative to the number of brainstem nuclei, PLS regression was used to derive latent components that maximize the covariance between regional gene expression profiles and brainstem-to-cortical weighted degree alterations. The first PLS component (PLS1), representing a weighted linear combination of gene expression profiles, was retained to capture the gene expression profile most significantly associated with the spatial distribution of weighted degree alterations. Functional enrichment analysis was subsequently performed to interpret the biological significance of the PLS1-associated gene expression patterns. Genes were ranked according to their PLS1 weighting coefficients, and enrichment of Gene Ontology (GO) terms and Kyoto Encyclopedia of Genes and Genomes (KEGG) pathways was assessed using the Gorilla toolkit (http://cbl-gorilla.cs.technion.ac.il/) and the clusterProfiler package (https://github.com/YuLab-SMU/clusterProfiler) implemented in R (version 4.5.0, R Foundation for Statistical Computing, Vienna, Austria), respectively.

### Cross-field-strength validation

To independently validate the brainstem-cortical functional architecture identified at 7 Tesla and assess its reproducibility at conventional field strength, we performed a cross-field-strength validation using an independent 3-Tesla resting-state fMRI cohort from the Parkinson’s Progression Markers Initiative (PPMI; https://www.ppmi-info.org/). The PPMI cohort included 30 OA (age: 67.14 ± 11.83 years; 11 females) and 30 individuals with iRBD (age: 69.06 ± 5.13 years; 6 females). Importantly, these participants were independent of the 7-Tesla discovery cohort, providing an external dataset in which to test the generalizability of the observed brainstem-cortical connectivity patterns. The 3-Tesla fMRI data were preprocessed using the same pipeline as the 7-Tesla fMRI data, except that susceptibility-induced distortions were corrected using pairs of anterior-to-posterior (AP) and posterior-to-anterior (PA) phase-encoding images. FC matrices were subsequently constructed using the same parcellation and connectivity procedures as for the 7-Tesla fMRI data. Correspondence between 7-Tesla and 3-Tesla brainstem-cortical FC patterns was quantified by Spearman’s correlation across corresponding brainstem and cortical regions, separately for OA and iRBD. These analyses tested whether the spatial organization of brainstem-cortical connectivity observed with ultra-high-field imaging could be reproduced using conventional 3-Tesla fMRI.

### Statistical inference and multiple comparisons correction

Clinical assessments in individuals with iRBD were evaluated using one-sample *t*-tests against the corresponding normative cutoff values, with statistical significance set at *p* < 0.05. One-sample *t*-tests against zero were used to assess whether FC-distance correlations differed significantly from zero within each group, whereas paired-sample *t*-tests were used to compare within-brainstem FC with brainstem–cortical FC within participants. Group differences in demographics and imaging measures were assessed using Fisher’s exact test for categorical variables (e.g., sex) and one-way analysis of variance (ANOVA) for continuous variables (e.g., age, FC, FC-distance correlation, weighted degree, and cortex-to-brainstem connectivity gradient scores). Significant main effects of group were followed by post hoc pairwise comparisons. The primary contrasts of interest were OA versus YA, reflecting aging-related alterations, and iRBD versus OA, reflecting alterations associated with prodromal synucleinopathy beyond those observed in aging. Multiple comparisons were corrected using the Benjamini-Hochberg false discovery rate (FDR), with significance set at FDR-corrected *p* < 0.05. The statistical significance of individual predictor contributions in the cortical dominance analyses was assessed using spin permutation testing (*n* = 5000) to account for spatial autocorrelation. The regional pattern of altered cortex-to-brainstem weighted degree was spatially rotated on the spherical cortical surface, and the dominance analysis was recomputed for each spin to generate spatially constrained null distributions of predictor contributions. The observed contribution of each predictor was evaluated with its corresponding null distribution to determine statistical significance. Spin tests were performed separately for the aging and iRBD contrasts and for each predictor domain (neurophysiological, structural and metabolic, and chemoarchitectural domains). The statistical significance of brainstem functional-domain contributions and spatial transcriptomic associations was assessed using permutation testing (*n* = 5000). For the functional-domain analyses, the binarized significance maps of brainstem-to-cortical weighted degree alterations and brainstem-to-cortical weighted degree computed from FC with unimodal and transmodal cortical regions were randomly permuted across brainstem nuclei, and dominance analysis was recomputed for each permutation to generate null distributions of individual predictor contributions. For the spatial transcriptomic analyses, the spatial correlation between PLS1 scores and the *t*-statistic maps of brainstem-to-cortical weighted degree differences was assessed by randomly permuting the *t*-statistic values across brainstem nuclei and recomputing the correlation for each permutation to generate a null distribution of correlation coefficients. Observed functional-domain contributions and correlations between PLS1 scores and altered brainstem-to-cortical weighted degree were evaluated against their respective null distributions. Permutation tests were performed separately for the aging and iRBD contrasts, with each functional-domain predictor evaluated separately in the dominance analyses. Functional enrichment of GO terms and KEGG pathways was considered significant at FDR-corrected *p* < 0.05.

## Data availability

All connectivity matrices are available from the corresponding author upon reasonable request and will be made publicly available upon acceptance of the manuscript for publication. Neurophysiological, structural, metabolic and neurotransmitter receptor/transporter data are available in neuromaps (https://github.com/netneurolab/neuromaps). Transcriptomic data are available from the Allen Human Brain Atlas (AHBA; https://human.brain-map.org/). Independent 3-Tesla resting-state fMRI data for OA and iRBD were obtained from the Parkinson’s Progression Markers Initiative (PPMI; https://www.ppmi-info.org/). The Brainstem Navigator atlas labels are available in NITRC (https://www.nitrc.org/projects/brainstemnavig).

## Code availability

All code used to perform the analyses is available from the corresponding author upon reasonable request and will be made publicly available upon acceptance of the manuscript for publication. MATLAB version R2017a and R version 4.5.0 were used for running scripts.

## Acknowledgements

This work was funded by grants from the National Institute of Aging, National Institutes of Health (R01-AG063982) and the Michael J. Fox Foundation (MJFF-022672). The funders had no role in study design, data collection and analysis, decision to publish or preparation of the paper.

## Author contributions

L.H., K.S. and M.B. conceived the study. L.H. and M.B. wrote the paper. K.S., M.G.G-G., F.F.H., S.K., A.S., S.G., A.V. and M.B. collected the data. K.S. and S.C. ran the preprocessing of YA. K.S. and L.H. ran the preprocessing of OA and iRBD. M.B. assisted with data pre-processing. L.H. performed the formal connectivity analysis from preprocessed data, as well as the characterization of the functional connectivity patterns. K.S., M.G.G-G., F.F.H., S.C., S.K., A.S., S.G., and A.V. assisted with interpretation and helped to revise the paper. M.B. was the project administrator.

## Additional information

Supplementary information is available in supplementary materials.

