## Supplementary Materials for "Charting vulnerability of brainstem functional pathways in aging and prodromal synucleinopathy"

**Anatomical coregistration**

Two anatomical coregistration strategies were implemented to accommodate the sequential expansion of the study cohort while improving computational efficiency (Figure S1). For the initial cohort, comprising 20 young adults (YA; s01-s20), 11 older adults (OA; s01-s11), and 12 individuals with isolated rapid eye movement sleep behavior disorder (iRBD; s01-s12), a group-based optimal template was constructed by iteratively registering and averaging individual MEMPRAGE images using the Advanced Normalization Tools (ANTs, version 2.2.0; <https://github.com/ANTsX/ANTs>). The group-based optimal template was then registered to the MNI152NLin6Asym 1-mm isotropic template space using an affine transformation and a nonlinear warp. The transformation matrices from individual MEMPRAGE images to the optimal template and from the optimal template to the MNI space were concatenated to generate the full coregistration matrix used to align each MEMPRAGE image into MNI space (Figure S1A). Then, six additional OA participants (s12-s17) and four additional individuals with iRBD (s13-s16) were subsequently included. These participants underwent the same anatomical preprocessing procedures as the initial cohort. To improve computational efficiency without reconstructing the group-based template, their MEMPRAGE images were directly registered to the MNI152NLin6Asym 1-mm isotropic template on a subject-by-subject basis. Nonlinear spatial normalization was performed using an ANTs-based registration workflow combining affine and diffeomorphic transformations with parameter settings optimized for accurate alignment of brainstem anatomy (Figure S1B).

**Subcortical extension of brainstem connectivity alterations**

To determine whether the brainstem connectivity alterations observed with the cortex extend to subcortical and spinal structures, subject-specific FC matrices were expanded to incorporate 27 additional regions spanning the subcortex/diencephalon and spinal cord. Specifically, these regions included 4 bilateral diencephalic nuclei from the Brainstem Navigator (the bilateral lateral and medial geniculate nuclei, subthalamic subregions 1 and 2), the hypothalamus^1^, and the ventral and dorsal spinal cord^2^, together with 8 bilateral FreeSurfer-derived subcortical structures (caudate, putamen, pallidum, basal forebrain, thalamus, amygdala, hippocampus, and cerebellum; Figure S4A)^3^. Brainstem-subcortical FC and weighted degree showed a progressive decrease from YA to OA, and further to iRBD (one-way ANOVA; *F*_(2,50)_ = 10.67, *p* < 0.001; Figure S4B and S4C). Statistically significant reductions were found in the bilateral thalamus, right putamen, bilateral lateral and left medial geniculate nuclei, and dorsal spinal cord in aging, as well as bilateral hippocampus, bilateral pallidum, bilateral thalamus, left putamen, bilateral subthalamic nuclei, and bilateral lateral and medial geniculate nuclei in iRBD (FDR-corrected *ps* < 0.05; Figure S4D).

Subcortical structures, positioned between the brainstem and cortex, indicate that brainstem dysfunction in aging and iRBD may involve not only direct cortical connections but also distributed subcortical pathways^4^. Consistent with the brainstem-cortical findings, progressive reductions in brainstem-subcortical connectivity further indicate disruptions across multiple levels of brainstem-subcortical-cortical organization, with aging and prodromal synucleinopathy affecting distinct subcortical pathways. The thalamus represents an integrative hub linking ascending brainstem circuits with distributed cortical networks, and age-related changes in thalamic structure and connectivity have been consistently associated with declines in attention, information processing and memory^5^^,^^6^. Reduced brainstem connectivity with the lateral and medial geniculate nuclei further extends age-related brainstem alterations to visual and auditory pathways^7^. In contrast, iRBD-related brainstem-subcortical alterations are consistent with growing evidence that prodromal synucleinopathy affects subcortical networks beyond the brainstem, including basal ganglia, cortico-striatal, limbic and sensory circuits^8-10^. Previous resting-state studies have identified altered substantia nigra-putamen connectivity before overt motor impairment^11^, reduced cortico-striatal connectivity in individuals with RBD and subtle motor dysfunction^12^, and decreased pallidal connectivity within the basal ganglia network^10^. Pallidum and subthalamic involvement therefore provide a functional link between brainstem alterations and basal ganglia circuits central to subsequent parkinsonian manifestations, whereas hippocampal involvement extends vulnerability to limbic circuits relevant to non-motor symptoms of synucleinopathy. Our findings demonstrate that brainstem dysfunction in iRBD is already embedded within subcortical circuits before the emergence of clinically overt parkinsonism.

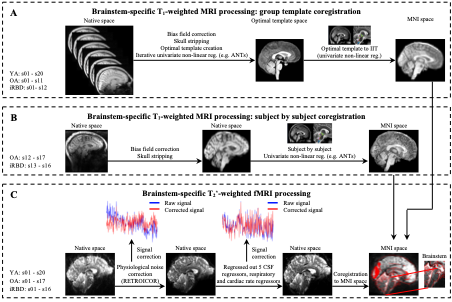

**Figure S1. Preprocessing of 7-Tesla resting-state fMRI data.** (A) Construction of group-based optimal anatomical templates. MEMPRAGE images from YA (s01-s20), OA (s01-s11) and iRBD (s01-s12) were bias-field corrected, skull-stripped, and iteratively registered using ANTs to generate group-based optimal templates, which were subsequently aligned to the MNI152 standard space through affine and nonlinear transformations. (B) Subject-by-subject preprocessing for additional participants (OA: s12-s17; iRBD: s13-s16). MEMPRAGE images were preprocessed with the same pipeline but directly registered to MNI152 standard space without group-based optimal anatomical template construction. (C) Brainstem-specific fMRI preprocessing workflow. Functional data underwent slice-timing correction, motion correction, fieldmap-based distortion correction, and RETROICOR correction of physiological noise. To minimize resampling, motion correction, distortion correction, and nonlinear coregistration to MEMPRAGE were applied in a single interpolation step. Brainstem-specific denoising included regression of six rigid-body motion parameters, RVT, HR, and five CSF-derived PCA components from the CSF spaces near the brainstem. The denoised fMRI time series were normalized to percent signal change, band-pass filtered (0.01–0.1 Hz), and the residual mean signal was removed.

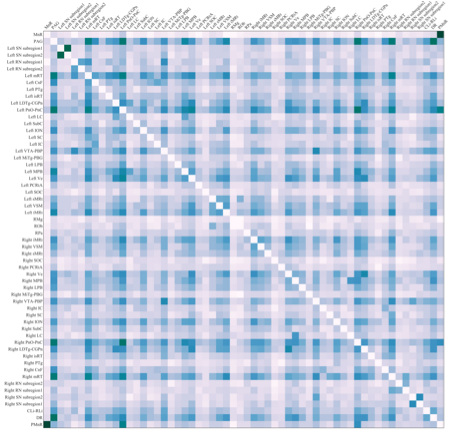

**Figure S2. Zoomed view of the brainstem-brainstem connectivity matrix with labels of brainstem nuclei.** The group-averaged brainstem-brainstem FC matrix in YA is shown as an example. Brainstem nuclei were delineated using the Brainstem Navigator (version 1.1; <https://www.nitrc.org/projects/brainstemnavig>). The atlas comprises 58 brainstem regions (8 midline and 25 bilateral nuclei).

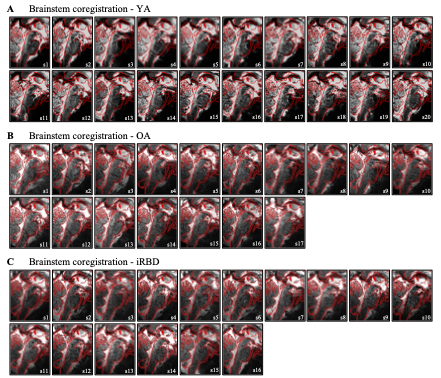

**Figure S3. Spatial alignment of fMRI to MNI space.** Coregistration accuracy is shown for YA (A), OA (B), and iRBD (C). For each subject, the time-averaged, motion-corrected fMRI volume from the first run (used as the reference for alignment) was underlaid (grayscale). The edges of the MNI152 standard template were overlaid in red to enable visual assessment of registration quality. Quality-control images were generated for all participants on the medial sagittal slice (MNI, x = 0).

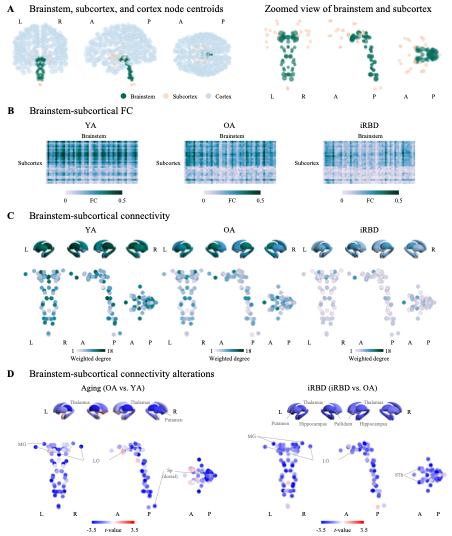

**Figure S4. Extension to subcortical and diencephalic structures.** FC construction was extended to the 8 bilateral FreeSurfer subcortical, 4 bilateral diencephalic and 3 brainstem-adjacent structures from the Brainstem Navigator. (A) For each brain region, the centroid coordinate is plotted with colors indicating structure. Grey nodes: 400 neocortical structures; orange nodes: 16 FreeSurfer and 11 Brainstem Navigator subcortical structures; green nodes: 58 Brainstem Navigator brainstem nuclei. (B) Brainstem-subcortical FC matrices across groups. Values represent Fisher z-transformed correlation coefficients. Color bar denotes FC strength. (C) FreeSurfer subcortical and Brainstem Navigator brainstem and diencephalic centroid coordinates sized and colored by volume size and weighted degree, reflecting how much each subcortical and diencephalic nucleus is connected with the 58 brainstem structures. (D) Group differences in weighted degree of brainstem-subcortical FC in aging (left) and iRBD (right). Node and structure colors represent the *t* values of weighted degree differences, and significant regions are labeled.

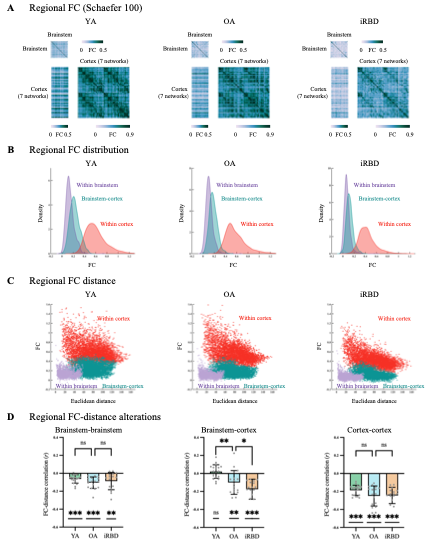

**Figure S5. Replication of FC using 100 cortical regions.** Main findings were repeated with the Schaefer 100-region cortical parcellation. (A) Regional FC matrices in YA, OA, and iRBD. Color bars denote FC strength for brainstem-brainstem, brainstem-cortical, and cortico-cortical connections. (B) FC density distributions across groups, shown separately for within-brainstem (purple), brainstem-cortical (green), and cortico-cortical (red) connections. (C) Scatter plots of FC between regions as a function of Euclidean distance between parcel centroids within the brainstem, between brainstem and cortex, and within the cortex across groups. (D) Group-level comparisons of FC-distance relationships across groups. Error bars represent the standard deviation (SD). * *p* < 0.05, ** *p* < 0.01, *** *p* < 0.001, *ns* not significant.

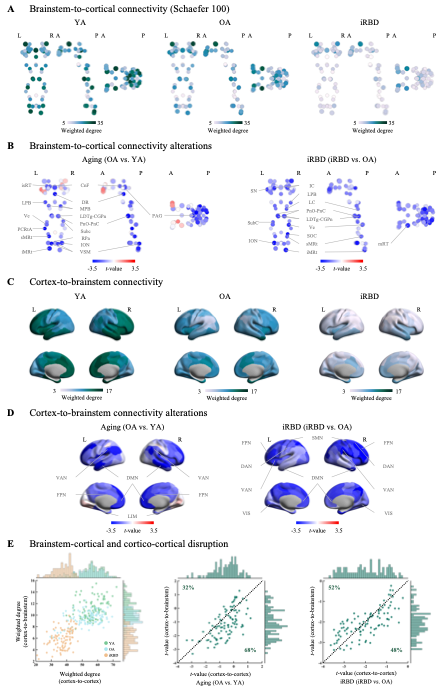

**Figure S6. Replication of weighted degree using 100 cortical regions.** Findings of weighted degree were repeated with the Schaefer 100-region cortical parcellation. (A) Brainstem-to-cortical weighted degree in YA, OA, and iRBD, with node size and color indicating volume size and weighted degree, respectively. (B) Group differences in brainstem-to-cortical weighted degree in aging and iRBD. Color bars denote *t* values of weighted degree differences. (C) Cortex-to-brainstem weighted degree in YA, OA, and iRBD, with color bars representing weighted degree. (D) Group differences in cortex-to-brainstem weighted degree in aging and iRBD. Color bars denote the *t* values of weighted degree differences. (E) Correspondence between cortex-to-cortex and cortex-to-brainstem weighted degree across groups (left). Spatial correspondence between regional *t* values of cortex-to-cortex and cortex-to-brainstem weighted degree changes in aging (middle) and iRBD (right). Percentages indicate the proportions of cortical regions lying above and below the identity line, reflecting relatively greater cortex-to-cortex and cortex-to-brainstem weighted degree changes, respectively.

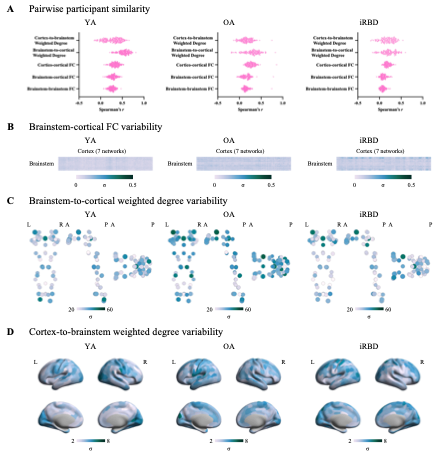

**Figure S7. Individual variability in FC and weighted degree.** (A) Pairwise correlations (Spearman’s *r*) between participants across groups, computed for brainstem-to-cortical weighted degree (YA: *r* = 0.54 ± 0.114, OA: *r* = 0.29 ± 0.177, iRBD: *r* = 0.22 ± 0.155; shown in Figure 2A), cortex-to-brainstem weighted degree (YA: *r* = 0.28 ± 0.181, OA: *r* = 0.16 ± 0.127, iRBD: *r* = 0.19 ± 0.148; shown in Figure 2C), and all FC (YA: *r* = 0.26 ± 0.066, OA: *r* = 0.16 ± 0.075, iRBD: *r* = 0.08 ± 0.05 for brainstem-brainstem FC; YA: *r* = 0.28 ± 0.075, OA: *r* = 0.16 ± 0.101, iRBD: *r* = 0.09 ± 0.063 for brainstem-cortical FC; YA: *r* = 0.33 ± 0.083, OA: *r* = 0.27 ± 0.094, iRBD: *r* = 0.19 ± 0.062 for cortico-cortical FC). (B) Inter-individual variability of brainstem-cortical FC, shown as SD across participants across groups. (C-D) Standard deviation of brainstem-to-cortical (C) and cortex-to-brainstem (D) weighted degree across participants within each group. Color bars denote SD of weighted degree.

**Table S1. Demographic characteristics of iRBD patients.**

| **iRBD**  **Subject ID** | **Sex**  **(M/F)** | **Age**  **(year)** | **RBDSQ** | **PSQI** | **ESS** | **MOCA** | **MDS-UPDRS Part III** | **Medication**  **(mg)** |
| --- | --- | --- | --- | --- | --- | --- | --- | --- |
| s1 | F | 68 | 9 | 6 | 4 | 26 | 0 | Melatonin (5) |
| s2 | M | 70 | 9 | 4 | 3 | 26 | 0 | None |
| s3 | M | 71 | 11 | 6 | 4 | 26 | 0 | None |
| s4 | M | 72 | 9 | 7 | 3 | 28 | 0 | Melatonin (6) |
| s5 | M | 66 | 10 | - | 8 | 27 | 0 | None |
| s6 | M | 71 | 9 | 8 | 3 | 27 | 0 | Melatonin (5) |
| s7 | M | 68 | 8 | 6 | 3 | 27 | 0 | Clonazepam (0.5) |
| s8 | M | 75 | 11 | 5 | 1 | 26 | 1 | Melatonin (9)/ Clonazepam (0.25) |
| s9 | M | 61 | 8 | 9 | 5 | 30 | 3 | None |
| s10 | M | 75 | 7 | 3 | 2 | 25 | 3 | None |
| s11 | M | 62 | 13 | 7 | 5 | 24 | 13 | Melatonin (5) |
| s12 | M | 56 | 11 | 4 | 13 | 30 | 9 | None |
| s13 | F | 60 | 10 | 8 | 6 | 28 | 2 | None |
| s14 | M | 75 | 7 | 6 | 8 | 27 | 16 | Melatonin (10)/ Clonazepam (0.5) |
| s15 | M | 63 | 10 | - | 10 | 25 | 0 | Melatonin (8) |
| s16 | F | 62 | 11 | - | 9 | 30 | 1 | Melatonin (5) |
| Summary | 13M/3F | 67.19 ± 5.96 | 9.56 ± 1.63 | 6.08±1.75 | 5.44±3.31 | 27±1.83 | 3±5.073 |  |
| T |  |  | 11.18 | 2.21 | 0.53 | 2.19 |  |  |
| df |  |  | 15 | 12 | 15 | 15 |  |  |
| *p* |  |  | < 0.001 | 0.023 | 0.302 | 0.022 |  |  |
| Cutoff value |  |  | >5 RBD episodes | >5 poor sleep quality | >5 abnormal daytime sleepiness | >26 normal cognition |  |  |

Note: Data are presented as mean ± standard deviation (SD). RBDSQ = REM Sleep Behavior Disorder Screening Questionnaire, PSQI = Pittsburgh Sleep Quality Index, ESS = Epworth Sleepiness Scale, MOCA = Montreal Cognitive Assessment, MDS-UPDRS = Movement Disorder Society sponsored revision of the Unified Parkinson’s Disease Rating Scale. The average RBDSQ score across individuals with iRBD was significantly higher than the normative cutoff value (*t*_(15)_ = 11.18, *p* < 0.001). Similarly, the average PSQI score was significantly higher than the normative cutoff (*t*_(12)_ = 2.21, *p* = 0.023). In contrast, the average ESS score did not differ significantly from the normative cutoff (*t*_(15)_ = 0.53, *p* = 0.302), whereas the average MoCA score was significantly higher than the normative cutoff (*t*_(15)_ = 2.19, *p* = 0.022), indicating no evidence of abnormal daytime sleepiness or global cognitive impairment in iRBD.

**Table S2. Functional classification of brainstem nuclei based on literature review.**

| **Nucleus** | **Sensory** | **Motor** | **Autonomic** | **Arousal** | **Limbic** | **Sleep** | **Cognitive** |
| --- | --- | --- | --- | --- | --- | --- | --- |
| Cuneiform nucleus (CnF) | 0 | 1^13^ | 0 | 1^2^ | 0 | 1^14^ | 0 |
| Dorsal raphe nucleus (DR) | 1^15^ | 0 | 1^15^ | 1^16^ | 1^17^ | 1^2^ | 0 |
| Inferior colliculus (IC) | 1^7^ | 0 | 0 | 1^18^ | 0 | 0 | 0 |
| Periaqueductal gray (PAG) | 0 | 0 | 1^16^ | 1^2^ | 1^15^ | 1^2^ | 1^19^ |
| Red nucleus - subregion 1 (RN1) | 0 | 1^16^ | 0 | 0 | 0 | 0 | 0 |
| Red nucleus - subregion 2 (RN2) | 0 | 1^16^ | 0 | 0 | 0 | 0 | 0 |
| Superior colliculus (SC) | 1^7^ | 1^15^ | 0 | 0 | 0 | 0 | 1^20^ |
| Substantia nigra, compatible with reticulata (SN1) | 0 | 1^16^ | 0 | 1^2^ | 1^21^ | 1^22^ | 1^23^ |
| Substantia nigra, compatible with compacta (SN2) | 0 | 1^16^ | 0 | 1^2^ | 1^21^ | 1^24^ | 1^23^ |
| Ventral tegmental area - parabrachial pigmented nucleus complex (VTA-PBP) | 0 | 1^25^ | 0 | 1^26^ | 1^25^ | 1^27^ | 1^15^ |
| Mesencephalic reticular formation (mRt) | 0 | 1^25^ | 0 | 1^26^ | 0 | 0 | 0 |
| Microcellular tegmental nucleus -  parabigeminal nucleus (MiTg-PBG) | 1^15^ | 0 | 0 | 0 | 0 | 0 | 0 |
| Isthmic reticular formation (isRt) | 0 | 1^26^ | 0 | 1^26^ | 0 | 0 | 0 |
| Locus coeruleus (LC) | 0 | 0 | 1^15^ | 1^2^ | 1^15^ | 1^2^ | 1^26^ |
| Laterodorsal tegmental nucleus - central gray of the rhombencephalon (LDTg-CGPn) | 0 | 0 | 0 | 1^2^ | 1^26^ | 1^2^ | 1^28^ |
| Lateral parabrachial nucleus (LPB) | 1^29^ | 0 | 1^29^ | 1^29^ | 1^30^ | 0 | 0 |
| Medial parabrachial nucleus (MPB) | 1^29^ | 0 | 1^29^ | 1^29^ | 1^30^ | 0 | 0 |
| Median raphe nucleus (MnR) | 0 | 0 | 0 | 1^16^ | 0 | 0 | 1^31^ |
| Paramedian raphe nucleus (PMnR) | 0 | 0 | 0 | 1^13^ | 0 | 0 | 0 |
| Pedunculotegmental nucleus (PTg) | 0 | 1^13^ | 0 | 1^13^ | 0 | 1^2^ | 1^26^ |
| Pontine reticular nucleus -  oral and caudal parts (PnO-PnC) | 0 | 1^13^ | 0 | 1^13^ | 0 | 1^2^ | 0 |
| Superior olivary complex (SOC) | 1^7^ | 0 | 0 | 0 | 0 | 0 | 0 |
| Subcoeruleus nucleus (SubC) | 0 | 1^26^ | 0 | 0 | 0 | 1^2^ | 0 |
| Vestibular nuclei complex (Ve) | 1^15^ | 1^15^ | 1^32^ | 1^15^ | 0 | 0 | 0 |
| Inferior olivary nucleus (ION) | 1^26^ | 1^16^ | 0 | 0 | 0 | 0 | 0 |
| Parvicellular reticular nucleus - alpha part (PCRtA) | 0 | 1^2^ | 1^15^ | 1^2^ | 1^15^ | 0 | 0 |
| Raphe magnus (RMg) | 0 | 0 | 1^16^ | 1^2^ | 1^15^ | 1^2^ | 0 |
| Raphe obscurus (ROb) | 0 | 1^2^ | 1^33^ | 0 | 1^15^ | 0 | 0 |
| Raphe pallidus (RPa) | 0 | 1^2^ | 1^34^ | 0 | 0 | 0 | 0 |
| Viscero-sensory-motor nuclei complex (VSM) | 1^29^ | 1^2^ | 1^29^ | 0 | 0 | 0 | 0 |
| Inferior medullary reticular formation (iMRt) | 0 | 1^2^ | 1^15^ | 1^2^ | 0 | 1^2^ | 0 |
| Superior medullary reticular formation (sMRt) | 0 | 1^2^ | 1^15^ | 1^2^ | 0 | 1^2^ | 0 |
| Caudal-rostral linear raphe (CLi-RLi) | 0 | 0 | 0 | 1^13^ | 1^26^ | 0 | 0 |

**Note:** A value of 1 indicates evidence in the literature that the brainstem nucleus participates in or modulates the corresponding functional domain. It does not imply that the domain represents the nucleus’s primary or exclusive function.
